# State-dependent signatures of Anti-NMDA Receptor Encephalitis: a dynamic functional connectivity study

**DOI:** 10.1101/2020.06.12.141945

**Authors:** Nina von Schwanenflug, Stephan Krohn, Josephine Heine, Friedemann Paul, Harald Prüss, Carsten Finke

**Author notes:** **Corresponding author:** Carsten Finke Charitéplatz 1, 10117 Berlin, Germany.

## Abstract

**Objective:** Traditional static functional connectivity (FC) analyses have shown functional network alterations in patients with anti-NMDA receptor encephalitis (NMDARE). Here, we use a dynamic FC approach that increases the temporal resolution of connectivity analyses from minutes to seconds. We hereby explore the spatiotemporal variability of large-scale brain network activity in NMDARE and assess the discriminatory power of functional brain states in a supervised classification approach.

**Methods:** We included resting-state fMRI data from 57 patients and 61 controls to extract four discrete connectivity states and assess state-wise group differences in FC, dwell time, transition frequency, fraction time and occurrence rate. Additionally, for each state, logistic regression models with embedded feature selection were trained to predict group status in a leave-one-out cross-validation scheme.

**Results:** Compared to controls, patients exhibited diverging dynamic FC patterns in three out of four states mainly encompassing the default-mode network and frontal areas. This was accompanied by a characteristic shift in the dwell time pattern and higher volatility of state transitions in patients. Moreover, dynamic FC measures were associated with disease severity, disease duration and positive and negative schizophrenia-like symptoms. Predictive power was highest in dynamic FC models and outperformed static analyses, reaching up to 78.6% classification accuracy.

**Conclusions:** By applying time-resolved analyses, we disentangle state-specific FC impairments and characteristic changes in temporal dynamics not detected in static analyses, offering new perspectives on functional reorganization underlying NMDARE. Correlation of dynamic FC measures with disease symptoms and severity indicates their clinical relevance and potential as prognostic biomarkers in NMDARE.

## INTRODUCTION

Anti-*N*-methyl-D-aspartate receptor encephalitis (NMDARE) is a severe autoimmune disorder of the central nervous system caused by antibodies targeting the NR1 subunit of the NMDA receptor.[1] The disease is characterized by a complex neuropsychiatric syndrome with delusions, hallucinations, movement abnormalities, autonomic dysfunction, decreased levels of consciousness, and cognitive dysfunction, e.g. deficits of executive control and memory.[1–5]

Despite the severe disease course, routine clinical magnetic resonance imaging (MRI) reveals no abnormalities in 50-80% of patients.[5,6] In contrast, functional connectivity (FC) is disrupted in distinct functional networks including medial-temporal, fronto-parietal, and visual networks.[7] Specifically, hippocampal connectivity with medial prefrontal regions of the default-mode network is significantly impaired, and these alterations are associated with the severity of memory impairment. Moreover, disruption of fronto-parietal and ventral attention networks correlates with positive and negative schizophrenia-like symptoms.[3,7] These traditional resting-state FC analyses have thus contributed to reveal the mechanisms underlying clinical symptoms in NMDARE by assessing the coherence of brain activity between distinct regions. However, traditional FC analyses are ‘static’ in the sense that blood-oxygen-level dependent (BOLD) time series are averaged across a scan with a duration of several minutes.

Yet, the brain is a complex dynamic system in which strength and spatial organization of connectivity patterns can change within seconds, resulting in multiple spatiotemporal organization patterns during one MRI scan.[8–10] ‘Dynamic’ FC approaches capture these changes of functional brain organization and allow for the investigation of temporal properties, i.e., identification of distinct connectivity states and analysis of transition trajectories between these states - alterations of which may vary with disease.[8] Indeed, recent studies report intriguing evidence that dynamic FC analyses enable a better characterization of network alterations in psychiatric and neurological diseases compared to static FC approaches.[9] Therefore, dynamic FC measures are increasingly suggested as novel biomarkers for disorders such as schizophrenia, major depression, stroke, and Alzheimer’s disease.[11–14]

One common method to analyse dynamic FC applies a clustering algorithm to obtain distinct functional brain states, which are defined as time-varying, but recurrent patterns of FC.[15] This approach provides a specifically promising tool for disentangling the dynamic network changes underlying the diverse neuropsychiatric symptoms in NMDARE. Here, we used this approach to (i) investigate the spatiotemporal properties of brain states in a large sample of patients with NMDARE and healthy controls (HC); (ii) explore the relationship between state dynamics, disease severity and duration, and psychiatric symptoms; and (iii) evaluate the potential of each brain state to discriminate between patients and controls using a supervised machine learning approach.

## METHODS

### Participants

For this study, 57 patients with NMDARE (female: 50, median age: 25.00 ± 14.50 years) were recruited from the Department of Neurology at Charité-Universitätsmedizin Berlin. Diagnosis was based on clinical presentation and detection of IgG NMDAR antibodies in the cerebrospinal fluid. Median disease duration, i.e. days spent in hospitalization, was 63 days (± 56.50, *N* = 45). Disease severity at the time of scan was assessed based on the modified Rankin scale (mRS; median mRS: 1.00 ± 1.00, *N* = 50). The control group consisted of 61 age- and sex-matched healthy participants (female: 54, median age: 26.00 ± 11.00 years) with no history of neurological or psychiatric disease. Clinical and demographic characteristics are summarized in supplementary table 1. All participants gave written informed consent, and the study was approved by the local ethics committee.

### MRI data acquisition

Structural and functional MRI data were acquired at the Berlin Center for Advanced Neuroimaging at Charité-Universitätsmedizin Berlin using a 20-channel head coil and a 3T Trim Trio scanner (Siemens, Erlangen, Germany). For resting-state functional MRI (rs-fMRI), we employed an echoplanar imaging sequence (TR=2.25s, TE=30ms, 260 volumes, voxel size=3.4×3.4×3.4mm^3^). High-resolution T1-weighted structural scans were collected using a magnetization-prepared rapid gradient echo sequence (MPRAGE; 1×1×1mm^3^).

### MRI data analysis

Our processing pipeline followed the procedure of recent related work.[15] Preprocessing of rs-fMRI scans included discarding the first 5 volumes to account for equilibration effects, slice time correction, realignment to the first volume, spatial normalization to MNI space (voxel size 2×2×2mm), and spatial smoothing with a 6 mm full width at half maximum smoothing kernel using the CONN Toolbox (https://web.conn-toolbox.org/).

#### Group independent component analysis

To perform group-independent component analysis (ICA) and dynamic functional network analysis, we applied the GroupICA fMRI toolbox (GIFT, http://mialab.mrn.org/software/gift/index.html). For each participant, 255 time points were first decomposed into 150 temporally independent principle components (PC), and subsequently into 100 independent PCs using the *Infomax* algorithm.[16] This procedure was repeated 20 times in ICASSO to estimate reliability and ensure stability of the decomposition.[17] For back-reconstruction of individual time courses and spatial maps, *gig-ica* (integrated in the GIFT Toolbox) was applied to the data.[18] The resulting 100 independent components were individually rated as signal or noise by three independent raters (NS, JH, CF). In total, 39 components were assigned to functional networks based on the labels proposed by Yeo *et al.*[19] For cerebellar and subcortical components, two distinct networks were added. This yielded a total of seven functional resting-state networks including sensorimotor (SM), visual (VIS), subcortical (SB), cerebellar (CB), default mode (DMN), dorsal attention (dATT), and frontoparietal control network (FPN). supplementary figure 1 shows all functional networks and supplementary table 2 contains peak values and coordinates for all components. Finally, we applied additional processing steps including linear, quadratic, and cubic detrending, motion regression (12 motion parameters) to reduce motion-related artifacts, high-frequency cut-off at 15Hz, despiking (identified as framewise displacement >0.5 mm), and interpolation of time courses using a 3rd order spline fit.

#### Static functional network connectivity analysis

To compare the dynamic FC results with conventional ‘static’ FC, we calculated the average pairwise connectivity between all component pairs across the resting-state scan using Pearson’s correlation coefficient *r* for each subject. Subsequently, age, sex, and motion parameters were regressed out, and Fisher z-transformation was applied.

#### Dynamic functional network connectivity analysis

In order to obtain FC dynamics, FC between all component pairs was calculated over consecutive windowed segments of the time courses (i.e., sliding windows) using a window of 30TR length (≙67.5 seconds) that shifted in steps of 1TR (≙2.25 seconds). After the correlation matrix was computed on each window (i.e., 225 39×39 matrices per participant), Fisher z-transformation was applied and age, sex, and motion parameters were regressed out as nuisance variables. Subsequently, matrices of each participant were concatenated, and k-means clustering was applied with k=4 according to the elbow criterion (see supplementary figure 2). Thus, each window was assigned to one of 4 clusters representing discrete network FC states.[15] Squared Euclidean distance was applied for clustering, and the process was repeated 100 times to avoid convergence on local minima.

#### Group differences in static and dynamic functional network connectivity (FNC)

For a global characterization of the static and the state-wise correlation matrices, modularity (as a measure of functional network segregation) and absolute mean connectivity (referred to as ‘overall connectivity’) were calculated.[20,21] In the static FC analysis, both measures were calculated on each subject’s connectivity matrix and subsequent group comparison was performed using a non-parametric t-test as applied in Glerean *et al*.[22] In the dynamic FC analysis, modularity and absolute mean connectivity were calculated for all windows in each state and averaged for each subject. Subsequently, a two-way ANOVA was conducted to estimate group- and state-wise effects as well as their interaction. For post-hoc analysis, a Kruskal-Wallis test was performed.

Next, we assessed group differences in FC between all component pairs for the static and the dynamic functional network analysis with respect to connectivity strength with a non-parametric t-test. For the dynamic FC analysis, group differences were evaluated for each state separately.

#### State dynamics

Besides the analysis of state-dependent connectivity patterns, estimation of time-varying FC provides the opportunity to capture dynamic metrics. Here, four commonly used metrics were calculated: (a) dwell time (i.e., average number of windows a participant spends in a particular state), (b) transition frequency (i.e., a participants’ number of transitions between each pair of states), (c) fraction time (i.e., percentage of windows spent in a state), and (d) state occurrence rate (i.e., number of participants that entered the state over the course of the scan).[11,14] Group differences in occurrence rates were estimated using the z-test for population proportions. For the other metrics, two-way ANOVAs were conducted to estimate group- and state-wise effects as well as their interaction. Post-hoc comparisons were evaluated with a non-parametric t-test or a Tukey’s test.

Between-group comparisons for the modularity and overall connectivity, static and dynamic functional network analysis, dwell time, fraction time, and occurrence rates were based only on participants that visited the respective state.

### State-wise classification

Finally, group-wise analyses were complemented by a supervised binary classification approach to assess the potential of the static FC markers and the four dynamic FC states to discriminate between patients and controls. As previous work has suggested visual, fronto-parietal, and default-mode network areas to represent the biologically relevant discriminatory features in NMDARE, these networks were considered as the set of input features.[7] For the static design and for each state, logistic regression models were trained on the z-scored FC indices to predict group status (NMDARE patients vs. HC) in a leave-one-out cross validation (LOOCV) scheme. To facilitate model sparsity and counteract overfitting, embedded feature selection was applied through L1 regularization. Hyperparameter optimization of the regularization strength λ was applied for each state input matrix (observations-by-connectivity features) by searching a linearly spaced parameter grid that was identical for all four states. Selection probability of each feature was read out as the empirical rate of non-zeroed feature weights over all predictions within a state. Prediction performance was evaluated by standard confusion matrix measures (i.e., true and false positive, and negative rates, and overall accuracy). Model training and prediction was implemented with the LIBSVM package for Matlab (The MathWorks, Inc., Natick, MA).

## RESULTS

### Functional Network Analysis

#### Static functional network connectivity analysis

We observed pairwise (component-to-component) differences in static FC between NMDARE patients and HC that clustered in the inter- and intra-connectivity of the DMN (figure 1 and table 1). In line with previous studies,[3,7] NMDARE patients showed decreased static connectivity between the hippocampus and the medial prefrontal cortex (mPFC; *p*_FDR_ < 0.05). In addition, NMDARE patients exhibited significantly reduced DMN connectivity with the supplementary motor area, TPO-junction, the parieto-occipital sulcus, and the superior frontal gyrus, and increased FC with the orbitofrontal gyrus (*p*_uncorr_ < 0.001). There was no significant difference between patients and controls in modularity (*mean* ± *SD*: 0.35 ± 0.09 vs. 0.33 ± 0.09; *t* = −1.19, *p* = 0.12) and overall connectivity (0.30 ± 0.05 vs. 0.31 ± 0.07; *t* = 0.63, *p* = 0.26).

**Table 1.** Test results of static functional network connectivity analysis. Table includes component name, network assignment, number (#), t-value, p-value and effect size (*d*) of component pairs that are highlighted in figure 1. * significant after FDR-correction.

| regions | network | component # | $p_{uncorr}$ | $p_{FDR}$ | T | d |
| --- | --- | --- | --- | --- | --- | --- |
| mPFC - Hippocampus | DMN – DMN | 36 – 59 | <0.0001* | 0.00024* | 4.36 | .62 |
| mPFC – SMA | DMN – SM | 36 – 23 | 0.00092 | 0.15 | 3.30 | .44 |
| mPFC – TPOJ | DMN – VIS | 33 – 38 | 0.00084 | 0.15 | 3.26 | .54 |
| mPFC – TPOJ | DMN – VIS | 36 – 38 | 0.00061 | 0.15 | 3.40 | .45 |
| mPFC – PHG | DMN – DMN | 36 – 14 | 0.00016 | 0.12 | 3.85 | .55 |
| SFG – POS | DMN – dATT | 24 – 80 | 0.00072 | 0.15 | 3.29 | .52 |
| mPFC – OFG | DMN – FPN | 33 – 29 | 0.00043 | 0.15 | -3.48 | -.37 |
| mPFC – SFG | DMN – DMN | 36 – 61 | 0.00087 | 0.15 | 3.27 | .40 |
| STG – SFG | DMN – FPN | 40 – 89 | 0.00057 | 0.15 | 3.24 | .49 |

**Figure 1.**
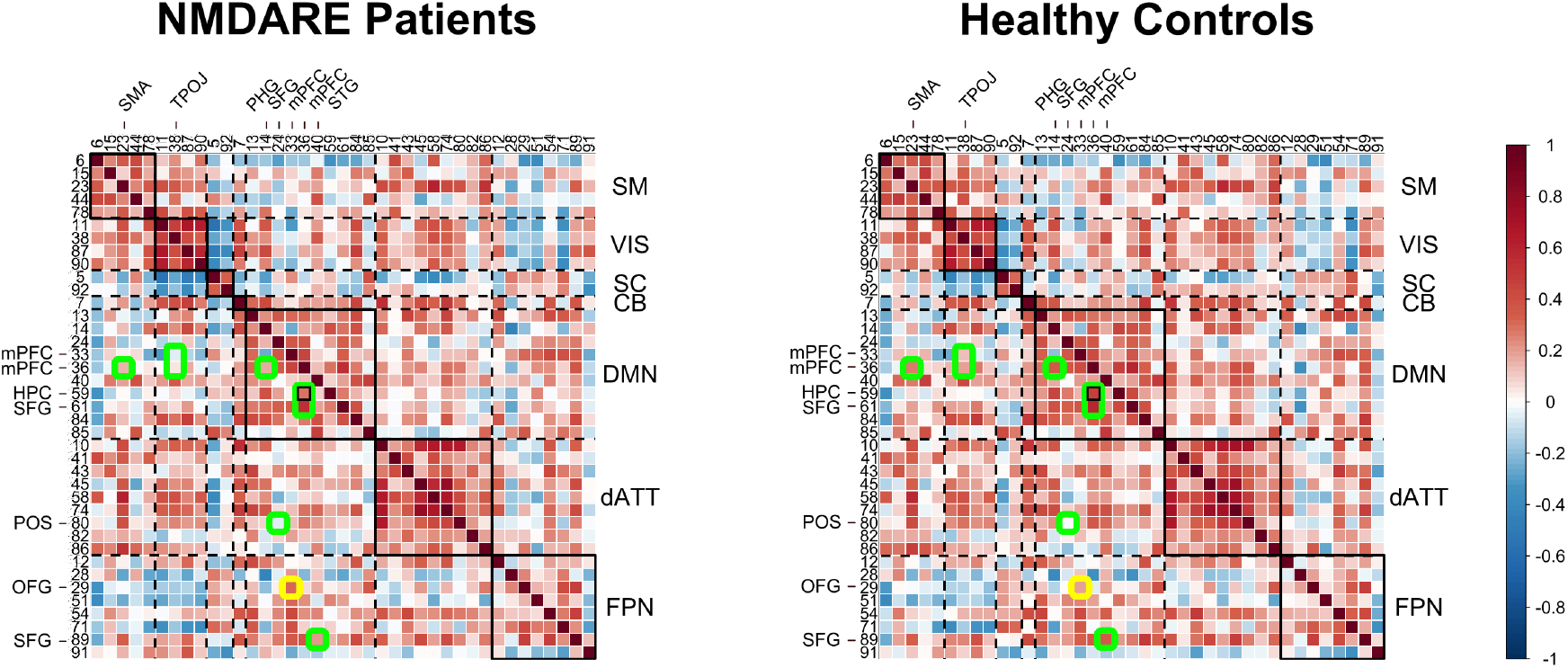
Mean static functional connectivity matrix across brain regions of NMDARE patients and healthy controls. Darker red/blue colors indicate higher positive/negative correlation values between component pairs. Green circles mark lower correlation values in NMDARE compared to controls and yellow circles indicate higher correlation values in NMDARE compared to controls. Small black rectangle indicates significant difference of FC between the hippocampus (region 59) and the medial prefrontal cortex (region 36) between patients and controls after FDR-correction (pFDR < 0.05), while no rectangle indicates differences between groups for puncorr < 0.001. Highlighted regions are displayed with anatomical labels. A key for the region numbers is provided in supplementary table 2. Big diagonal rectangles indicate functional networks, e.g. the sensorimotor network that is comprised of the regions 6, 15, 23, 44 and 78.

Following previous studies that found a correlation between the mPFC-hippocampal connection and disease severity variables,[3,7] we conducted a post-hoc correlation analysis (using Pearson’s correlation coefficient) between these regions and disease severity at the time of scan (mRS). Higher mRS scores were associated with a reduced connectivity between the parahippocampal gyrus and the mPFC (*r* = −0.33, *p* = 0.029), as well as with lower connectivity between the hippocampus and the mPFC (*r*= −0.25, *p* = 0.081), albeit limited to a statistical trend.

#### Dynamic functional network connectivity analysis

K-means clustering identified four connectivity states for HC and NMDARE patients. Group-wise mean connectivity states are shown in figure 3. Multiple regression models for modularity and overall connectivity yielded a significant effect for state (modularity, *p* < 0.001; overall connectivity, *p* < 0.001), but not for group or interaction. The dominant state 1 closely resembled the static FC pattern (*r* = 0.94, supplementary table 14) with low overall connectivity and moderate modularity. States 2 and 3 were both characterized by high overall connectivity, while only state 2 had a highly segregated structure (i.e., high modularity). In contrast to state 3, state 4 exhibited high modularity and low overall connectivity (figure 2, see supplementary tables 3–8 for detailed test statistics).

**Figure 2.**
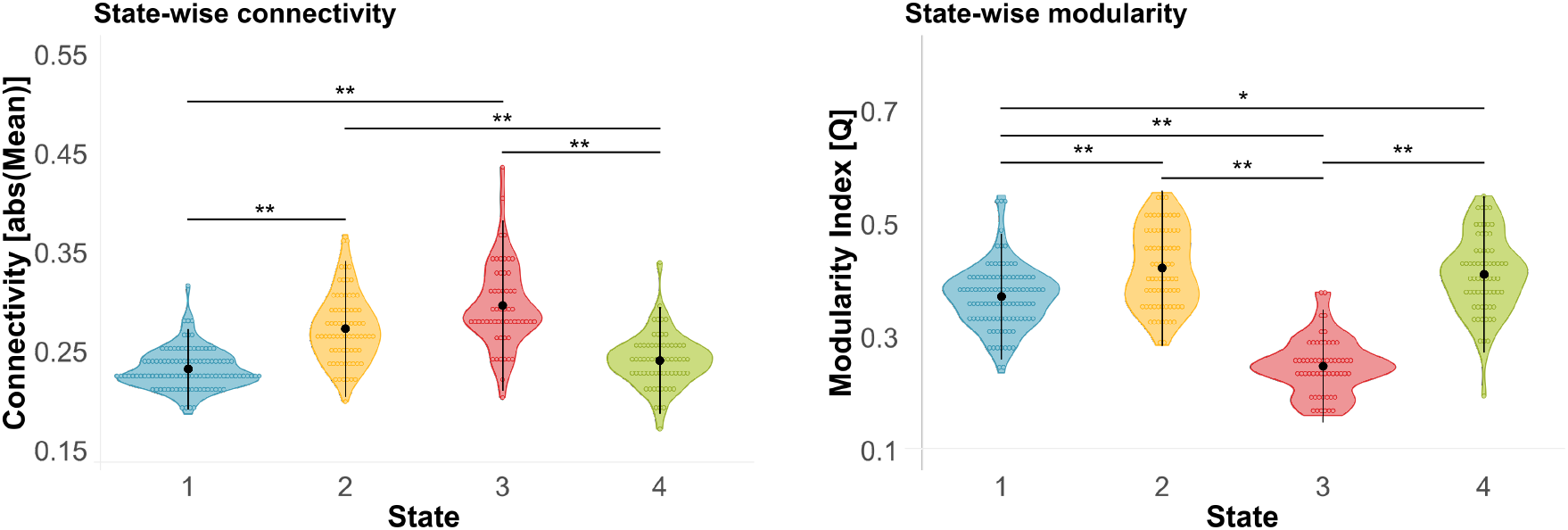
State-wise comparison of overall connectivity and modularity. In general, states 1 and 4 exhibited weak overall state connectivity compared to state 2 and 3. Segregation of functional networks, as measured with modularity, was highest in state 2 and 4, followed by state 1 and weakest in state 3. Black dots and vertical lines represent mean and standard deviation ** p < 0.001 (Bonferroni-corrected). * p < 0.01 (Bonferroni-corrected). Detailed test statistics can be found in supplementary tables 3–8.

**Figure 3.**
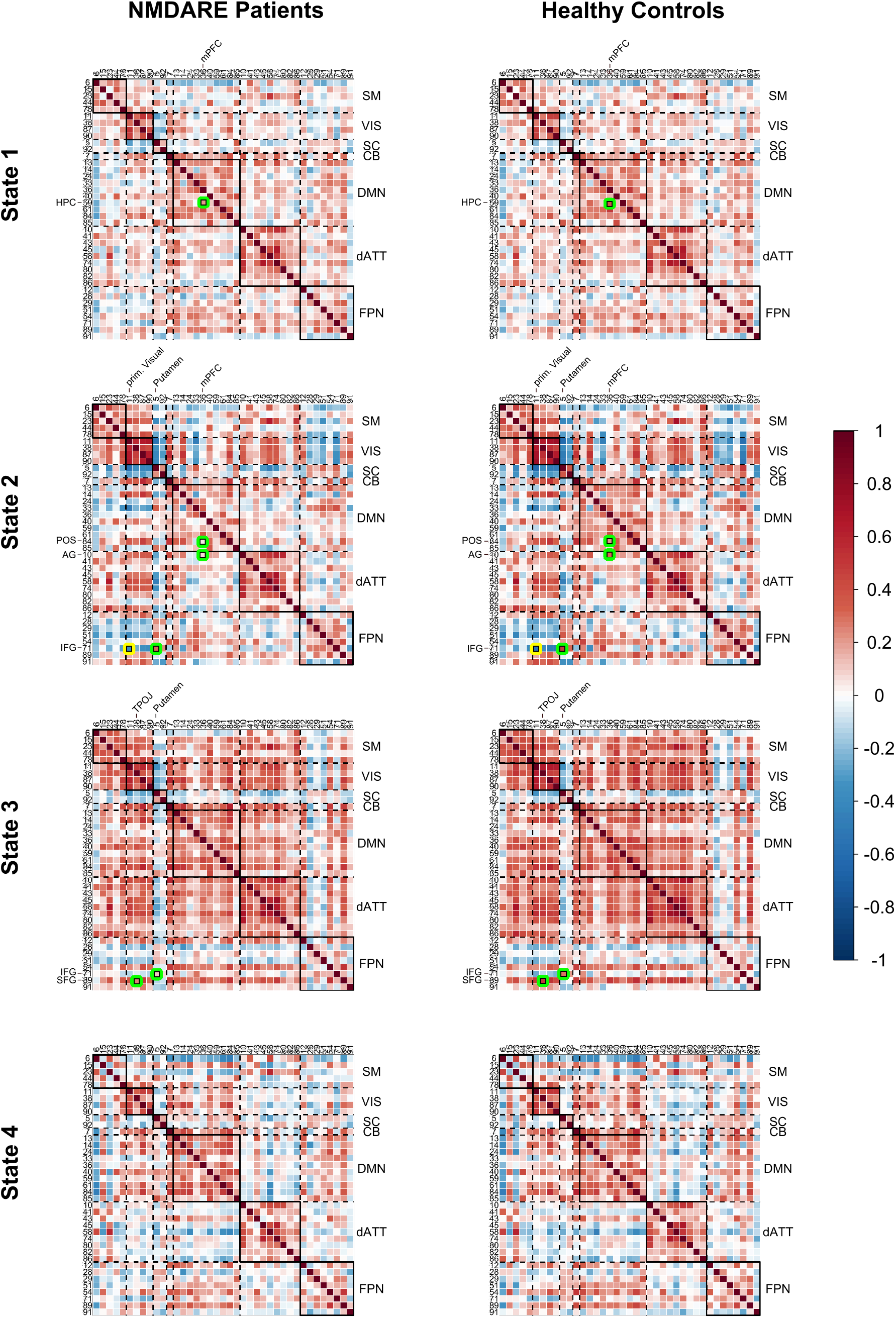
Dynamic functional network connectivity states for NMDARE patients and healthy controls. Darker red/blue colors indicate higher positive/negative correlation values between component pairs. Green circles mark lower correlation values in NMDARE compared to controls and yellow circles indicate higher correlation values in NMDARE compared to controls. Small black rectangles indicate significant differences of FC between patients and controls after FDR-correction (pFDR < 0.05). Highlighted regions are displayed with anatomical labels. A key for the region numbers is provided in supplementary table 2. Big diagonal black rectangles indicate functional networks, e.g. the sensorimotor network that is comprised of the regions 6, 15, 23, 44 and 78.

Anti-NMDA receptor encephalitis patients showed distinct FC alterations across the four connectivity states in comparison to controls (figure 3 and table 2). As in the static FC group analysis, group differences comprised the DMN, VIS and FPN, but in a state-dependent fashion: in the static FC-resembling state 1, patients with NMDARE displayed decreased connectivity between the mPFC and the hippocampus, i.e., results very similar to the findings in the static FC analysis. The highly modular state 2 showed impaired connectivity between the mPFC and the angular gyrus as well as the parieto-occipital sulcus in patients. Furthermore, the inferior frontal gyrus exhibited connectivity alterations with the putamen (bil.) and the visual cortex. Similarly, the densely-connected/highly-integrative state 3 was characterized by decreased connectivity from the inferior frontal gyrus to the putamen. Additionally, connectivity between the TPOJ and the superior frontal gyrus was reduced in NMDARE patients compared to HC. For state 4, no significant alterations were observed after FDR-correction.

**Table 2.**
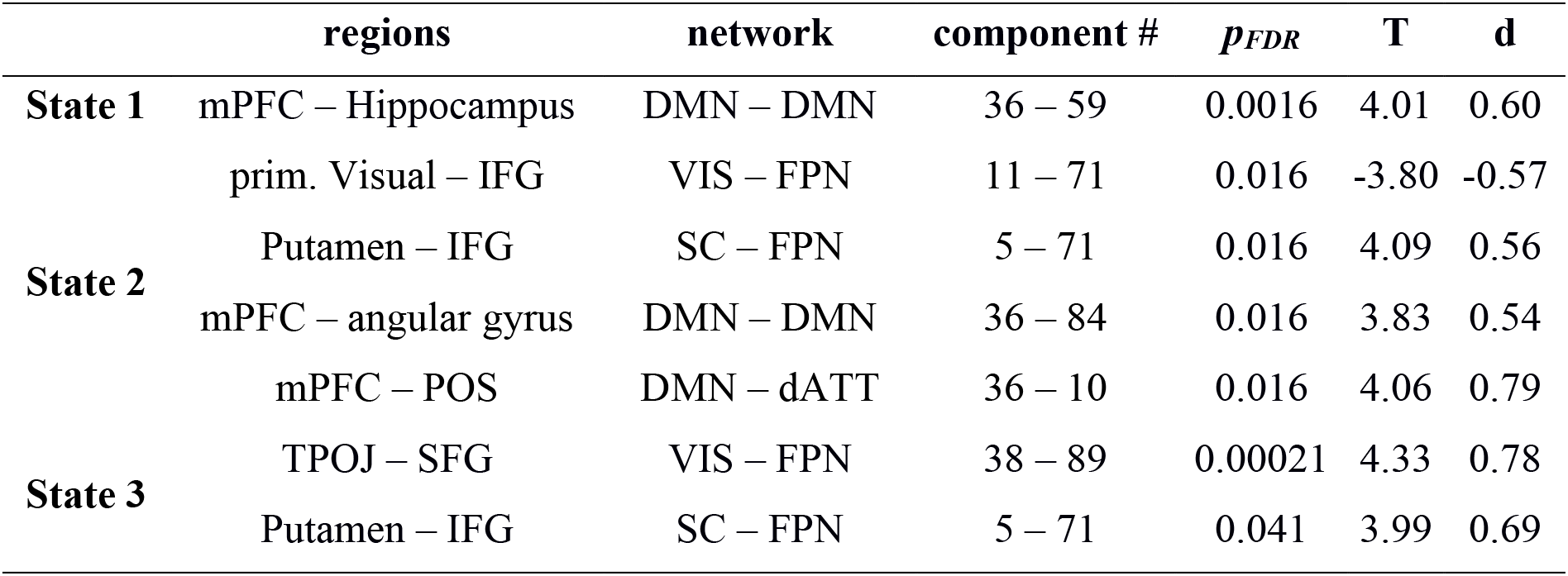
Test results of dynamic functional network connectivity analysis. Table includes component name, network assignment, number (#), t-value, p-value and effect size (*d*) of component pairs that are highlighted in figure 3.

Next, we obtained the correlation coefficient between all significant component pairs and disease severity (mRS at the time of scan) as well as disease duration (days in hospitalization): In the strongly segregated state 2, higher disease severity was significantly associated with a decrease in FC between mPFC and angular gyrus (*r* = −0.40, *p* = 0.013), while in the densely-connected/highly-integrative state 3, longer disease duration was significantly related to a decrease in connectivity between TPOJ and SFG (*r* = −0.60, *p* = 0.014). Due to the exploratory nature of the study, post-hoc correlation analyses were not corrected for multiple comparisons.

#### State dynamics

In addition to state-wise connectivity patterns, we assessed state and group differences in dwell time, transition frequency, fraction time, and occurrence rate using two-way ANOVAs. We found a significant state effect in *dwell times* (*p* = 0.00021): dwell times were higher for patients and controls in state 1 compared to state 2 (*T* = −3.77, *p* = 0.0010) and 3 (*T* = −3.61, *p* = 0.0019). Importantly, a significant group effect (*p* = 0.010) revealed a shift in dwell times between patients and controls: While patients showed lower dwell times in the dominant static FC-resembling state 1 (*p* = 0.020), they had higher dwell times in the strongly segregated state 2 compared to controls (*p* = 0.032; figure 4). Similarly, the model for *transition frequencies* yielded a significant effect for group (*p* = 0.044) and state (*p* < 0.001). Post-hoc group comparisons exhibited higher transition frequencies in patients between states with high and low overall connectivity, i.e., state 1 and 2 (*p* = 0.043), and between states with high and low across-network connectivity, i.e., states 3 and 4 (*p* = 0.0063; figure 4), in comparison to controls. Furthermore, transitions from/to state 1 were significantly more frequent than transitions from states 2/3 to state 4 or vice versa. *Fraction time* differed across states (*p* = 0.0023), but not between groups (*p* = 0.56). A post-hoc test revealed higher percentages of windows in state 1 compared to state 2 (*T* = −3.23, *p* = 0.0077) and 3 (*T* = −3.02, *p* = 0.014). *Occurrence rates* of dynamic FC states were similar in NMDARE patients and HC: The static FC-resembling state 1 showed the highest occurrence, followed by states 2 and 4; the lowest occurrence rates were observed for the densely-connected/highly-integrative state 3. Despite similar general occurrences, state-wise between-group proportion tests revealed that a higher number of patients visited the highly segregated state 2 compared to controls (*p* = 0.019), while the proportions were equal for both groups in states 1, 3 and 4. Detailed test-statistics can be found in table 3 and supplementary tables 9–13.

**Figure 4.**
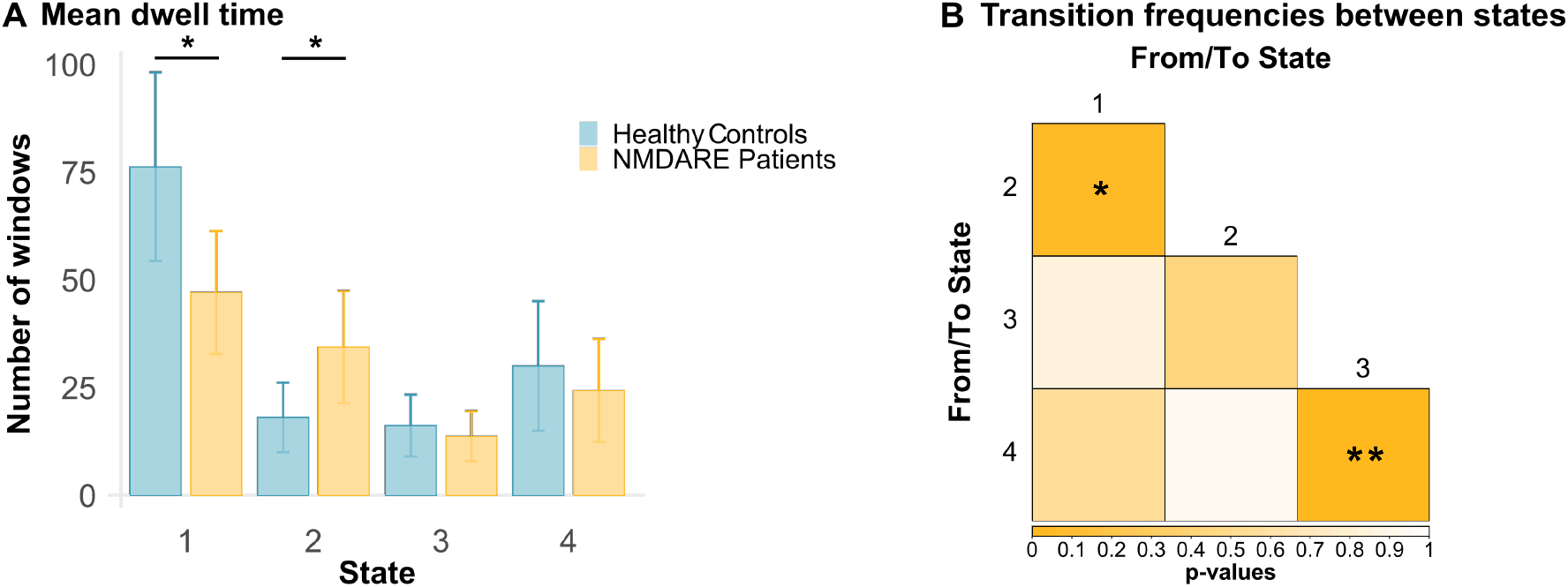
Group differences in state dynamics. (A) Group differences in average dwell time (in windows). Solid lines point to significant differences in post-hoc testing between groups (non-parametric t-test, uncorrected). (B) Group differences in transition frequencies between states (p-values). For transition frequencies, the direction of transition was ignored. Post-hoc group comparisons were calculated using a non-parametric t-test (uncorrected). * p < 0.05; ** p < 0.01.

**Table 3.**
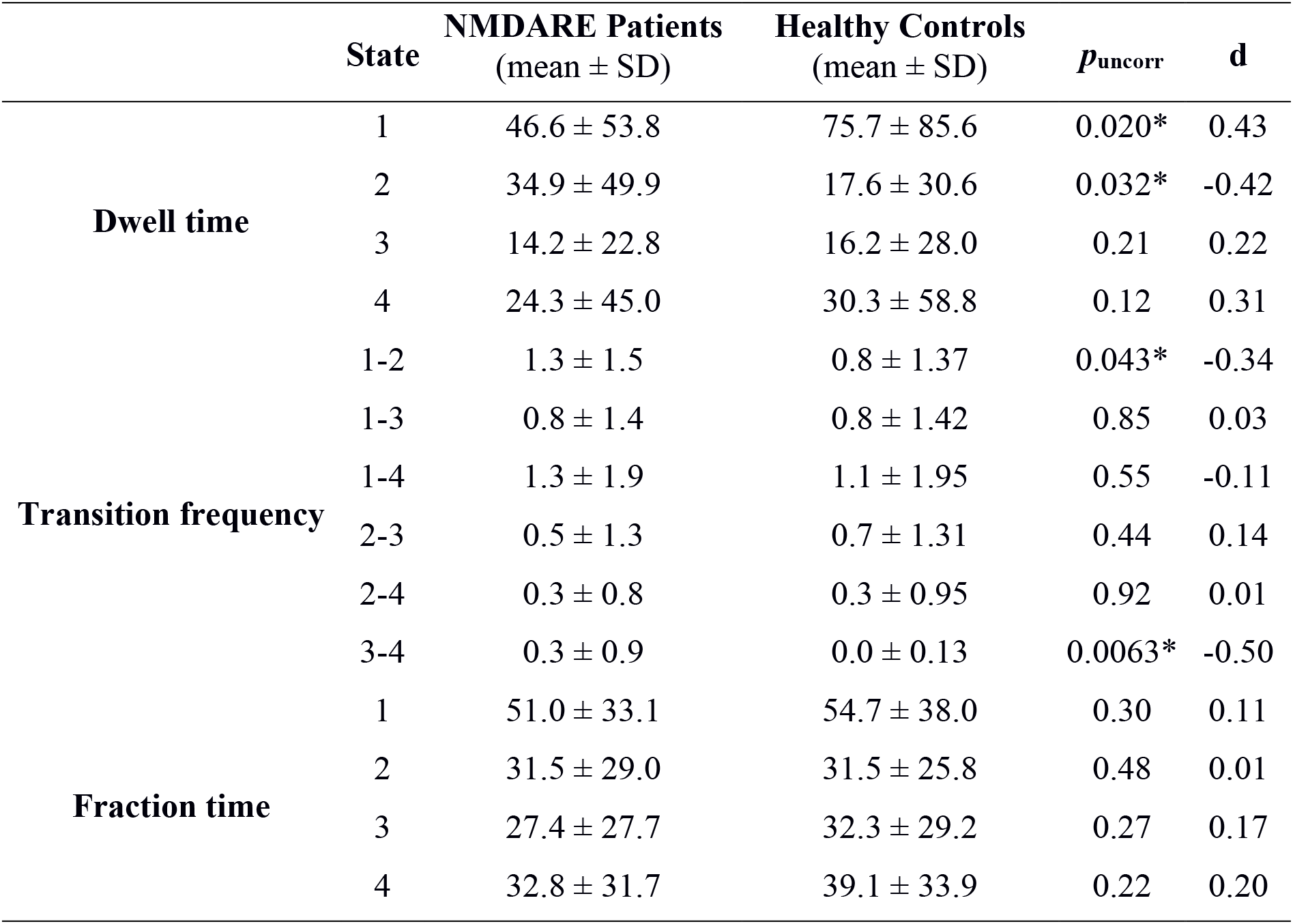
Group differences in dwell time (average number of windows), transition frequencies between states (absolute numbers), and fraction time (percentage). Group differences were calculated using a two-sided non-parametric t-test. p-values and effect sizes (*d*) are shown. * p < 0.05 (uncorrected).

| | State | NMDARE Patients<br>(mean ± SD) | Healthy Controls<br>(mean ± SD) | $p_{\text{uncorr}}$ | *d* |
| --- | --- | --- | --- | --- | --- |
| **Dwell time** | 1 | 46.6 ± 53.8 | 75.7 ± 85.6 | 0.020* | 0.43 |
| | 2 | 34.9 ± 49.9 | 17.6 ± 30.6 | 0.032* | -0.42 |
| | 3 | 14.2 ± 22.8 | 16.2 ± 28.0 | 0.21 | 0.22 |
| | 4 | 24.3 ± 45.0 | 30.3 ± 58.8 | 0.12 | 0.31 |
| **Transition frequency** | 1-2 | 1.3 ± 1.5 | 0.8 ± 1.37 | 0.043* | -0.34 |
| | 1-3 | 0.8 ± 1.4 | 0.8 ± 1.42 | 0.85 | 0.03 |
| | 1-4 | 1.3 ± 1.9 | 1.1 ± 1.95 | 0.55 | -0.11 |
| | 2-3 | 0.5 ± 1.3 | 0.7 ± 1.31 | 0.44 | 0.14 |
| | 2-4 | 0.3 ± 0.8 | 0.3 ± 0.95 | 0.92 | 0.01 |
| | 3-4 | 0.3 ± 0.9 | 0.0 ± 0.13 | 0.0063* | -0.50 |
| **Fraction time** | 1 | 51.0 ± 33.1 | 54.7 ± 38.0 | 0.30 | 0.11 |
| | 2 | 31.5 ± 29.0 | 31.5 ± 25.8 | 0.48 | 0.01 |
| | 3 | 27.4 ± 27.7 | 32.3 ± 29.2 | 0.27 | 0.17 |
| | 4 | 32.8 ± 31.7 | 39.1 ± 33.9 | 0.22 | 0.20 |

To identify a relationship between disease severity variables (i.e., acute days in hospitalization and mRS score at the time of scan) and dynamic metrics (i.e., dwell time and transition frequency), we conducted Pearson’s correlation analyses between these variables. We found that increased transition frequency between state 1 and 2 was associated with disease severity at the time of scan (*r* = 0.38, *p* = 0.0070), while increased transition frequency between state 3 and 4 was correlated with disease duration (days in hospitalization) in NMDARE patients (*r* = 0.37, *p* = 0.033). We further compared dwell time and transition frequency from patients with positive and negative schizophrenia-like psychiatric symptoms to those without respective psychiatric symptoms. Here, patients with positive symptoms exhibited higher dwell times (*z* = 2.33, *p* = 0.020) in the highly segregated state 4 compared to patients without positive symptoms. In contrast, patients with negative symptoms showed higher dwell times (*z* = 2.89, *p* = 0.0038) in the densely-connected/highly-integrative state 3 compared to those without negative symptoms.

### Classification analyses

Binary classification (NMDARE patients vs. HC) based on static connectivity features yielded an overall prediction accuracy of 72%, with balanced feature distribution across the networks (see supplementary figure 3). When dynamic connectivity features were considered, discriminatory power differed in a state-wise fashion. Prediction performance was lowest for the dominant, static FC-resembling state 1 (overall accuracy of 61.5%), intermediate and similar to model performance with static feature input for the modular-structured states 2 (72.6%) and 4 (70.8%), and highest for the least frequent and densely-connected/highly-integrative state 3 (78.6%; see supplementary figure 4 for the state-wise confusion matrices). Besides model evaluation outcomes, the feature selection frequencies over individual predictions in the LOOCV scheme also varied across states (figure 5). While states 1 and 3 yielded balanced selection rates over both across-and within-network connectivity features, states 2 and 4 showed fewer discriminatory features, and these were primarily across-network connections (FPN to VIS and DMN for state 2, and DMN to VIS for state 4). Importantly, although some connectivity features were discriminatory across several states (e.g., component pair 12-90 showed high selection frequency for states 1-3), the constellation of predictive features changed dynamically over connectivity states, emphasizing the uniqueness of each state.

**Figure 5.**
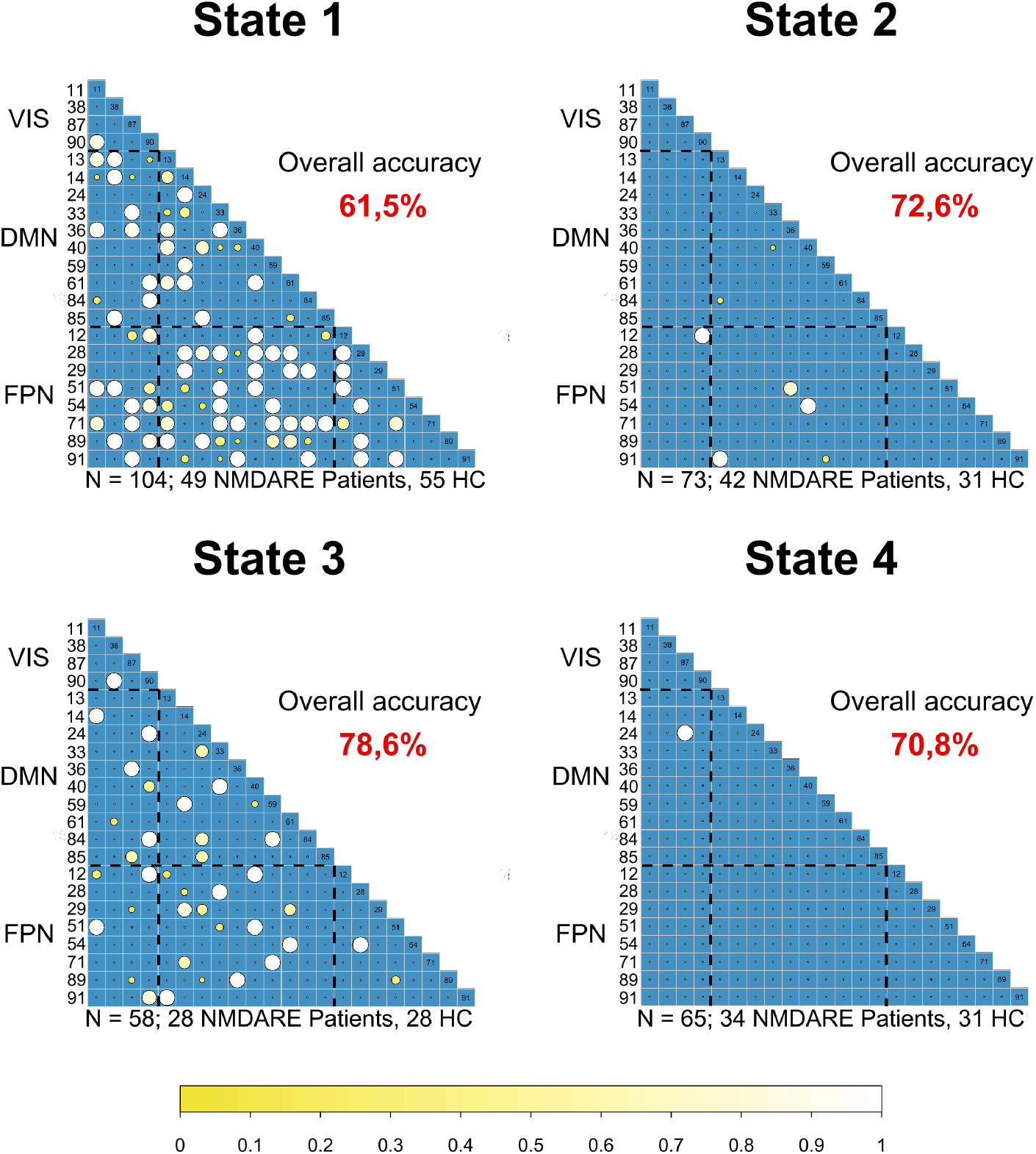
Feature selection matrices for state-wise predictions of group status (NMDARE patients vs healthy controls). The feature selection exceeding a minimum threshold at 10% of individuals within state predictions are displayed. Bigger and brighter circles indicate a higher selection rate. A key for the region numbers is provided in supplementary table 2.

## DISCUSSION

In this study, we applied dynamic functional connectivity analyses to characterize distinct connectivity patterns and temporal dynamics of network interactions in NMDARE. Investigating state-specific FC alterations, we found a marked impairment of functional connectivity between the hippocampus and the medial prefrontal cortex in the most visited, i.e., dominant state. This connectivity pattern closely mirrored observations in the static FC analysis and corroborated previous findings.[3,7] Three additionally identified states showed connectivity alterations within the default mode network and between frontal, visual, and subcortical areas – findings that remained undetected in the static FC analysis. Investigation of state dynamics showed a systematic shift in dwell time from the dominant state to a strongly segregated state in patients. Likewise, negative and positive schizophrenia-like symptoms were associated with distinct patterns of state preference. In addition, an increased volatility of transitions between states with high and low overall connectivity and states with high and low segregation was observed in patients. These state dynamics were associated with disease severity and duration. Finally, classification analyses revealed that discriminatory network features and predictive power varied dynamically across states, exceeding the discriminatory power of static FC analyses and yielding the highest prediction in a highly-connected/highly-integrated state. Our observations demonstrate the potential of time-resolved FC analysis for a better characterization of disease mechanisms involved in NMDARE and the value of dynamic FC measures as biomarkers for neuropsychiatric disorders.

### Static functional network connectivity analysis

In line with previous studies, conventional static FC analyses showed impaired connectivity between the mPFC and the hippocampus as well as altered connectivity patterns in frontal parts of the DMN.[3,7] Indeed, the CA1 subregion of the hippocampus and the prefrontal cortex contain the highest density of NMDA receptors.[23] Converging observations of disrupted hippocampal-prefrontal connectivity are thus biologically plausible and point to a robust disease biomarker and potential treatment target in NMDARE. Furthermore, both brain regions are main components of the DMN and are involved in memory and executive functions [24,25] – the two cognitive domains most frequently impaired in patients with NMDARE.[4,26,27]

### Dynamic functional network connectivity analysis

However, these findings are inherently limited to a static account of connectivity changes. Time-varying FC, in contrast, captures moment-to-moment changes in connectivity, reflecting a more physiologically plausible model of brain activity. One line of thought hypothesizes that the temporal variability of FC networks enables a systematic exploration of network configurations, which allows brain regions to dynamically (dis-)engage, and modulate changes in cognition and behavior.[28] Dynamic state analysis[15] as employed in this study, represents a powerful tool to describe these dynamics and potential instabilities of this process.

Indeed, state-wise group comparisons revealed connectivity differences between patients and controls in three out of four states. These differences were most pronounced in within- and across-network connectivity of the DMN and almost exclusively manifested as reduced connectivity strength in NMDARE.

State 1 represented the dominant state, i.e., the most visited state, the state in which participants remained longest and that was involved in most transitions. The connectivity pattern of state 1 was characterized by low overall connectivity and low segregation. Anti-NMDA receptor encephalitis patients showed a significantly impaired hippocampal-prefrontal connectivity in comparison to controls that closely resembled the pattern observed in current and previous static FC analyses.[3,7] Thus, the connectivity pattern in the dominant state 1 seems to drive findings of altered connectivity in conventional static FC analyses. In contrast, states 2 to 4 showed strikingly different features. FC alterations in states 2 and 3 went beyond the aggregated findings of the static analysis and revealed impaired connectivity between the mPFC and parieto-occipital areas, and between the IFG and the putamen (state 2). The latter is also present in state 3 along with impaired frontal-parietal connectivity.

Importantly, correlation analyses revealed that these dynamic FC alterations were associated with disease severity and disease duration, primarily involving mPFC connectivity and highlighting the clinical relevance of these findings. Together, these results disentangle state-specific connectivity patterns observed in conventional FC analyses and indicate the potential differential contribution of state-wise FC alterations to clinical symptoms and disease stages.

### State dynamics

In addition to these alterations in large-scale connectivity patterns in different states, NMDARE patients showed distinct temporal properties with respect to connectivity states, i.e., different transition frequencies and dwell times in comparison to controls. This involved a systematic shift in dwell time from the dominant state 1 to the segregated state 2, with patients nearly doubling their dwell time in state 2. Interestingly, recent evidence shows that successful working memory performance relies on increased network integration.[29] Prolonged dwelling in the segregated, less-integrated state 2 might thus be related to the frequently observed working memory deficits in NMDARE.[4] Remarkably, patients that experienced positive schizophrenia-like symptoms spent more time in the highly segregated state 4, while those with negative symptoms increased their dwell time in the highly integrative state 3. These observations are consistent with recent studies in schizophrenia showing an increased modular network structure in patients.[30]

Additionally, patients showed an increase in transition frequencies between states 1 and 2 as well as between states 3 and 4. These transition frequency alterations were significantly correlated with disease severity and duration, indicating that severe NMDARE disease courses are associated with more volatile transition dynamics, while state preference (i.e., dwell time) is not affected. The dynamic interplay between brain regions – in the sense of the flexible (dis-)engagement of functional links and state transitions – is critical to efficiently process internal and external stimuli and flexibly adapt behavior. While state transitions are thought to be generally important to explore different brain states in order to facilitate and enhance cognitive flexibility, overly unstable transition dynamics may be linked to deficiencies in the integration and stable representation of information.[28,31] The imbalance of stability and volatility may, therefore, lead to impaired memory, perception, or executive functions.[32,33] These suggestive links between state dynamics and impaired cognitive performance in NMDARE require further detailed investigations in combined task-based and resting state fMRI studies.

### Relation to schizophrenia

Our results show a notable convergence with recent studies in patients with schizophrenia reporting a similarly marked shift in state preference[11] as well as increased overall transition frequencies[34], and altered modular network structure.[30] Given the considerable overlap in psychiatric symptoms in patients with schizophrenia and NMDARE[35,36] and the glutamate hypothesis positing NMDA receptor dysfunction as pathophysiological basis for cognitive and psychiatric symptoms in schizophrenia,[37,38] our findings raise the interesting possibility that the transdiagnostic psychopathological profile of both diseases[39] could be paralleled by a common set of dynamic network alterations. This should be addressed in comparative studies of NMDARE and schizophrenia that use the same MRI acquisition and analysis protocols.

### Classification analyses

While our findings support the role of the hippocampus, the anterior DMN and frontal areas as potential connectivity biomarkers in NMDARE, group-level analyses are not suited to estimate the discriminatory power of connectivity alterations or their value to predict disease severity.[40] To this end, we applied classification analyses based on logistic regression models to these data. Prediction performance and the set of selected network features were variable across the different connectivity states, indicating that discriminatory network constellations differ between states. Interestingly, the best performance (78.6% accuracy) was achieved in state 3, which showed lowest overall occurrences but a highly integrative connectivity pattern. In contrast, static FC distinguished patients from controls with 72% accuracy. These results show that dynamic FC models can outperform static models and indicate the potential of spatiotemporal FC dynamics as prognostic biomarkers in NMDARE.

### Limitations

Some limitations of the present study deserve mentioning. First, window-based approaches require the specification of windowing parameters, and the optimal choices in this regard are an active area of research and debate.[41] Second, a given window size may only capture a part of the dynamic nature of the human brain, as networks may reconfigure over different time scales even within the possible temporal spectrum of MRI signals.[41] Lastly, for classification analyses, it is generally sensible to include large amounts of data.[40] While our study is based on a large study population in light of the incidence of the disease, the sample sizes per state varied as not all participants visited all states.

## Conclusions

Our analyses identified distinct brain states with characteristic patterns of FC alterations and shifted temporal dynamics in patients with NMDARE that remained undetected in conventional static analyses. Critically, dynamic FC measures correlated with disease severity and duration as well as with psychiatric symptoms, suggesting that altered resting-state dynamics carry relevant clinical information and emphasizing the potential of state-specific connectivity changes as biomarkers in NMDARE. Given converging findings in other neuropsychiatric diseases, time-resolved FC analysis holds promise for an improved characterization and understanding of brain functioning in these disorders.

## Contributors

NS designed the study, conducted the static and dynamic functional network connectivity analysis, led the writing of the manuscript and produced the figures. SK implemented the classification analysis. SK and CF (principal investigator) significantly contributed to the design & interpretation of the analysis and writing. NS, JH and CF rated the components from the group ICA. JH, FP, HP, and CF were involved in conceiving/conducting the cohort study and data collection. All authors edited and revised the manuscript.

## Supplemental material

**Supplementary Table 1:**
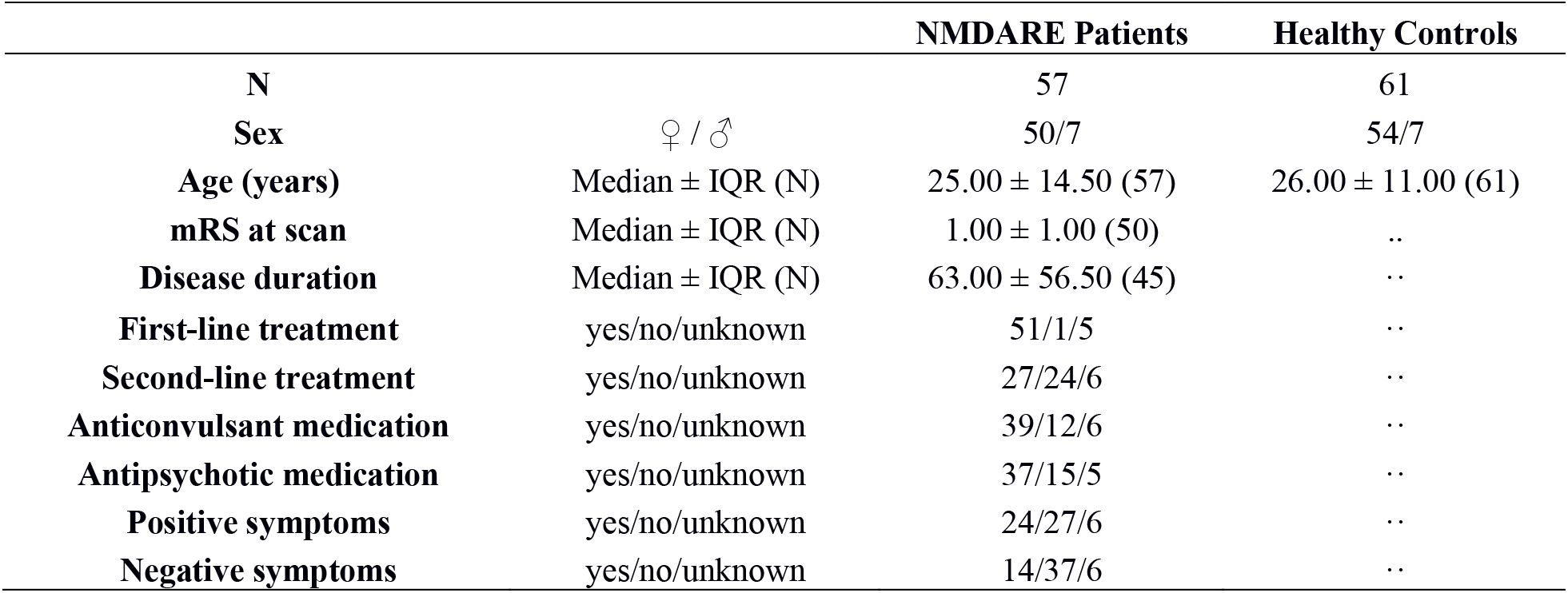
Demographic variables and clinical measures of the participants. Table lists median and interquartilerange (IQR) of age, mRS at scan, and disease duration. Treatment, medication, and psychiatric symptoms during disease course were evaluated using a binary (present: ‘yes’ vs. absent: ‘no’) scale. Disease duration = acute days in hospitalization; N = number of subjects.

**Supplementary Table 2:**
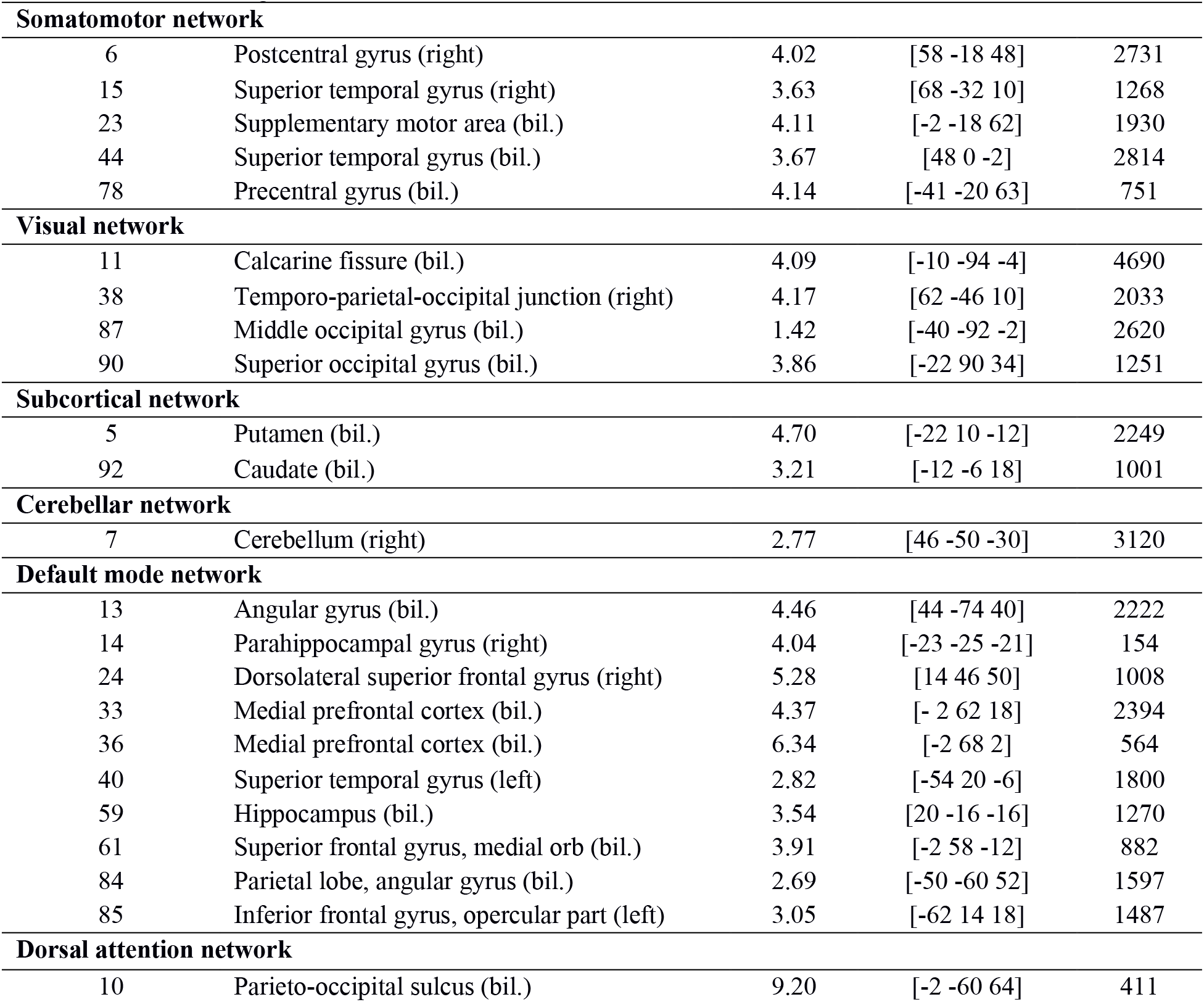
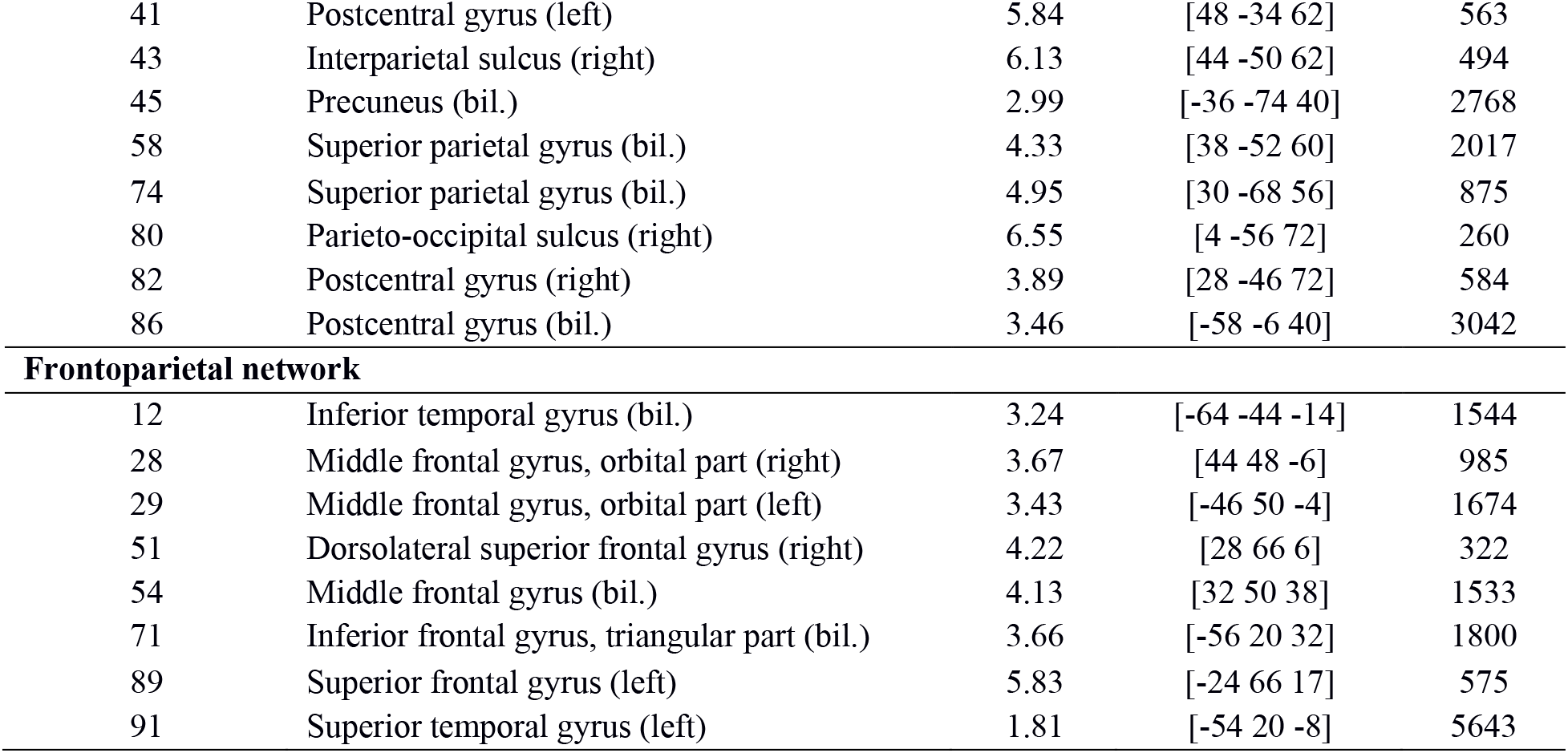
Location of included independent components. Component numbers, component labels, maximum t-value, MNI-coordinates of peak voxel and number of voxels in each component counting the voxels that contain the 60% highest values.

**Supplementary Table 3:**
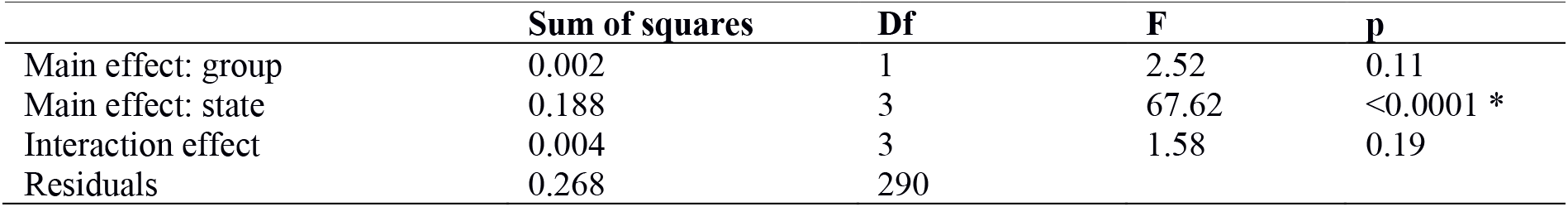
Two-way ANOVA for overall connectivity. * indicates significant effect.

**Supplementary Table 4:**
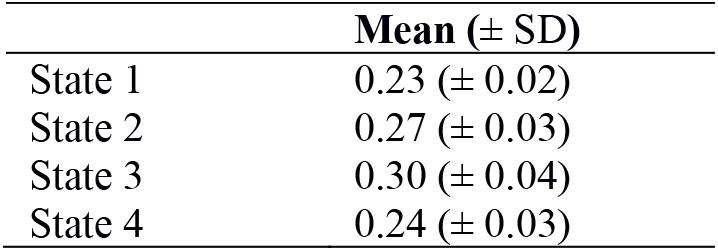
Average windows-wise overall connectivity (± SD) across all subjects.

| | Mean (± SD) |
| --- | --- |
| State 1 | 0.23 (± 0.02) |
| State 2 | 0.27 (± 0.03) |
| State 3 | 0.30 (± 0.04) |
| State 4 | 0.24 (± 0.03) |

**Supplementary Table 5:**
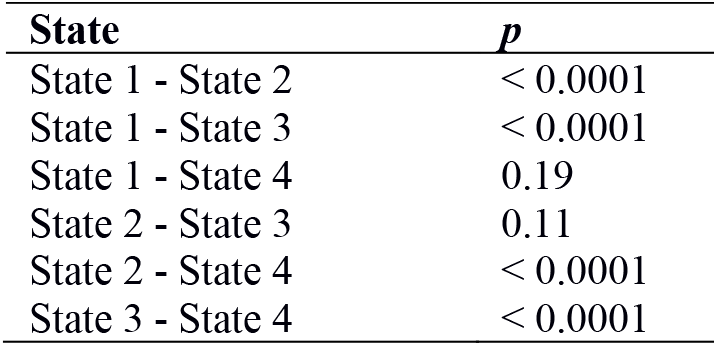
Post-hoc Kruskal-Wallis test to examine state-wise differences in overall connectivity (Chi^2^=124.37, p < 0.0001, df =3). The table contains the Bonferroni-corrected p-values for pairwise state comparison.

**Supplementary Table 6:**
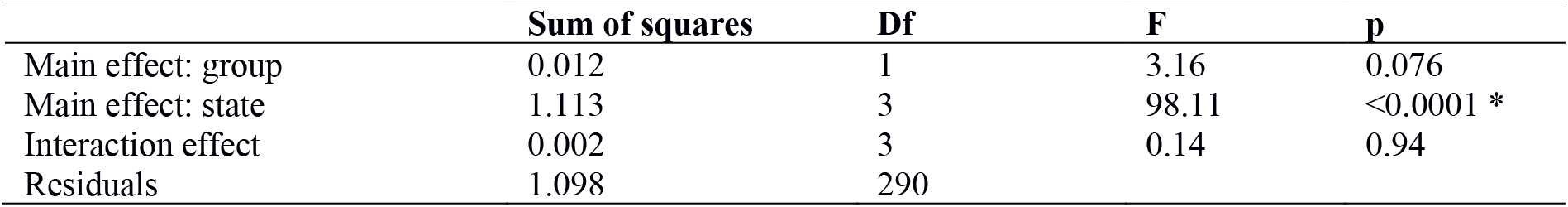
Two-way ANOVA for modularity. * indicates significant effect.

**Supplementary Table 7:**
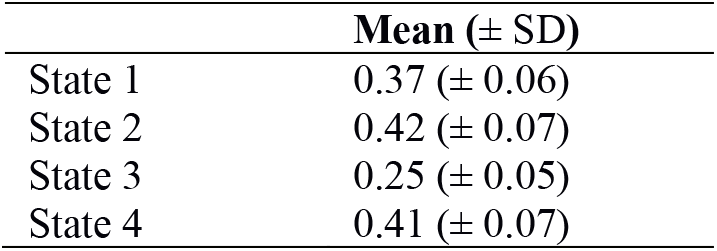
Average window-wise modularity (± SD) across all subjects.

| | Mean (± SD) |
| --- | --- |
| State 1 | 0.37 (± 0.06) |
| State 2 | 0.42 (± 0.07) |
| State 3 | 0.25 (± 0.05) |
| State 4 | 0.41 (± 0.07) |

**Supplementary Table 8:**
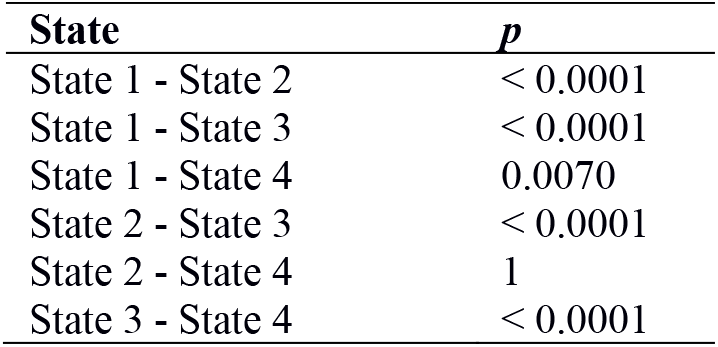
Post-hoc Kruskal-Wallis test to examine state-wise differences in modularity (Chi^2^=136.08, p < 0.0001, df =3). The table contains the Bonferroni-corrected p-values for pairwise state comparison.

**Supplementary Table 9:**
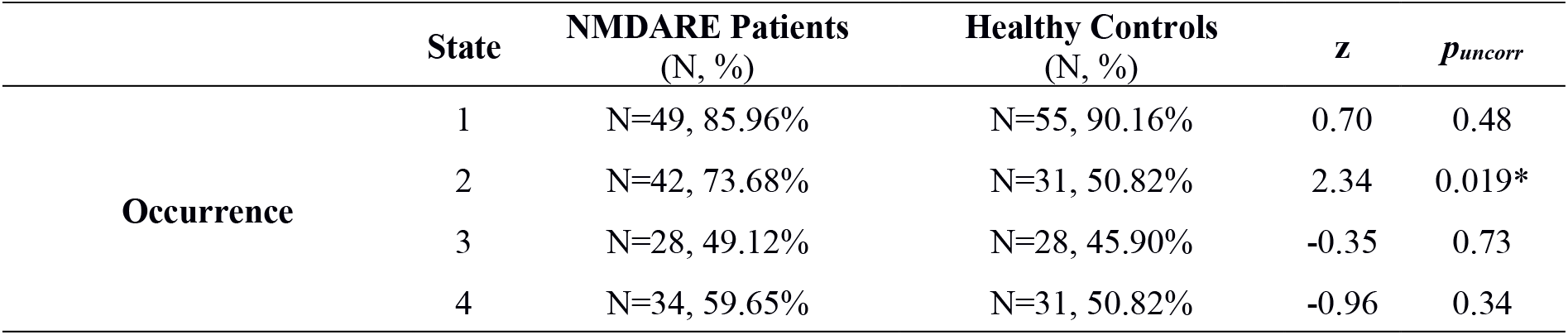
Group differences in occurrences of states. Group differences were calculated using the z-test for population proportions. * p < 0.05 (uncorrected).

**Supplementary Table 10:**
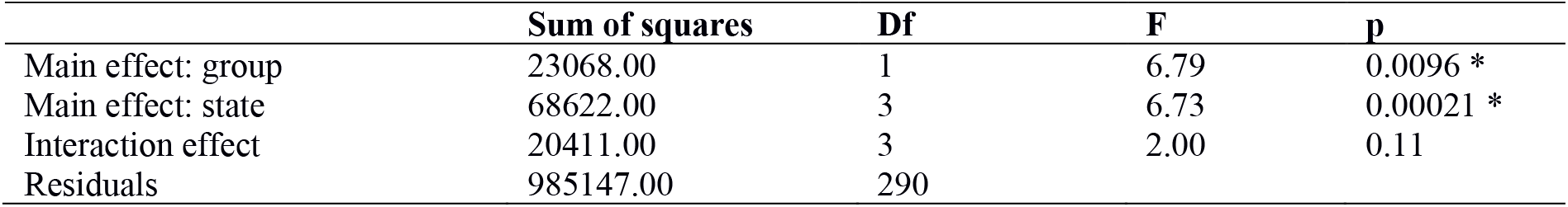
Two-way ANOVA for dwell time. * indicates significant effect.

**Supplementary Table 11:**
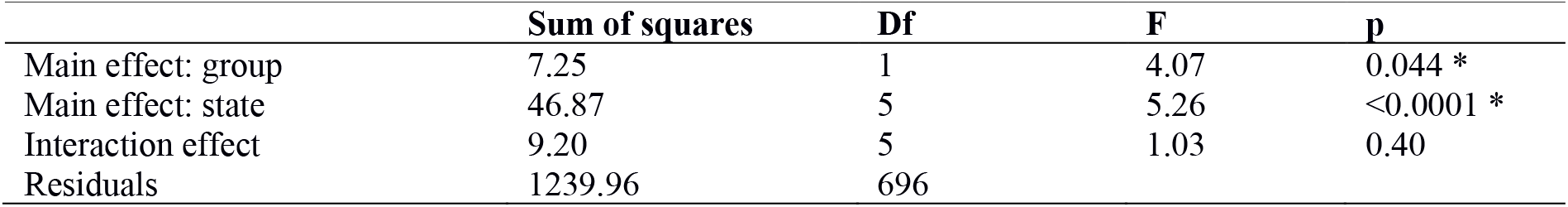
Two-way ANOVA for transition frequencies. * indicates significant effect.

**Supplementary Table 12:**
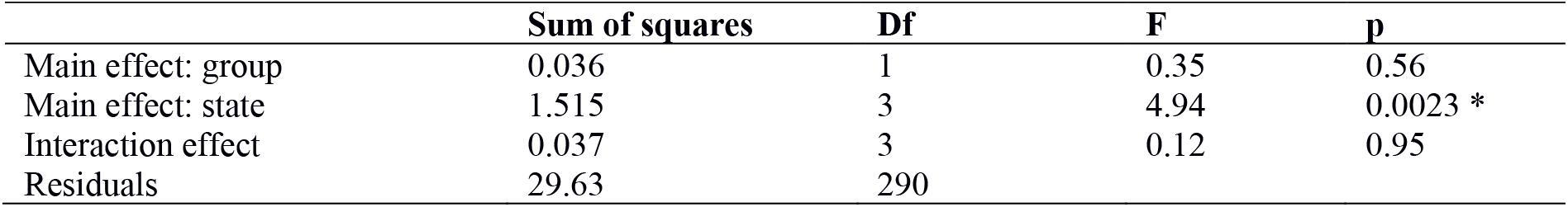
Two-way ANOVA for fraction time. * indicates significant effect.

**Supplementary Table 13:**
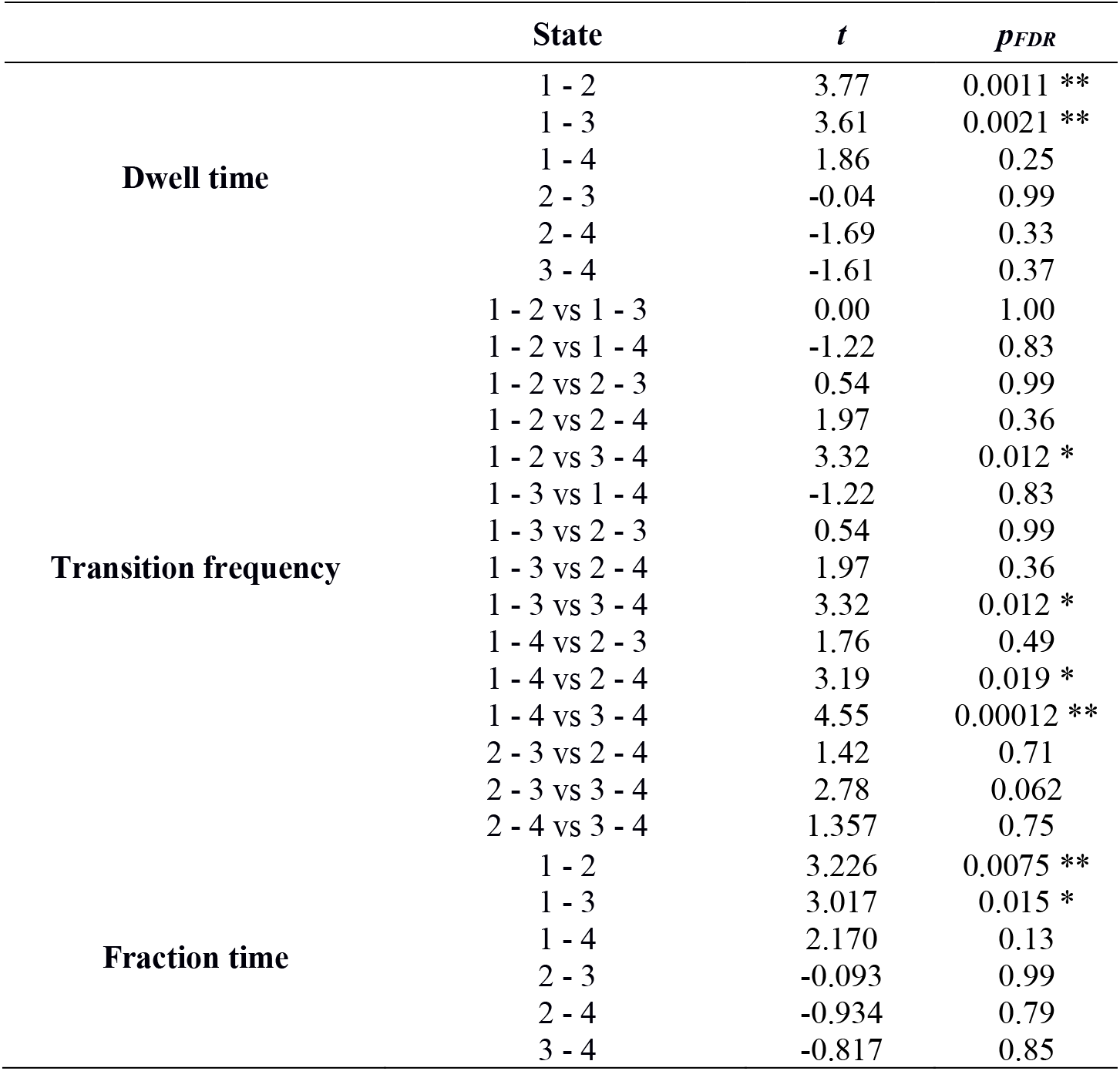
Differences between states in dwell time (windows), transition frequencies between states (absolute numbers), and fraction time (percentage). Differences between states were calculated using a Tukey’s test. T-values and p-values are shown. * p < 0.05 (FDR-corrected), ** p < 0.01 (FDR-corrected).

**Supplementary Table 14:**
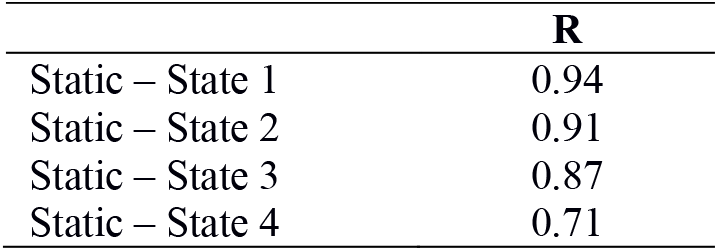
Pearson’s correlation coefficient between the participants’ average static FC and the participants’ average of each state.

**Supplementary Fig. 1:**
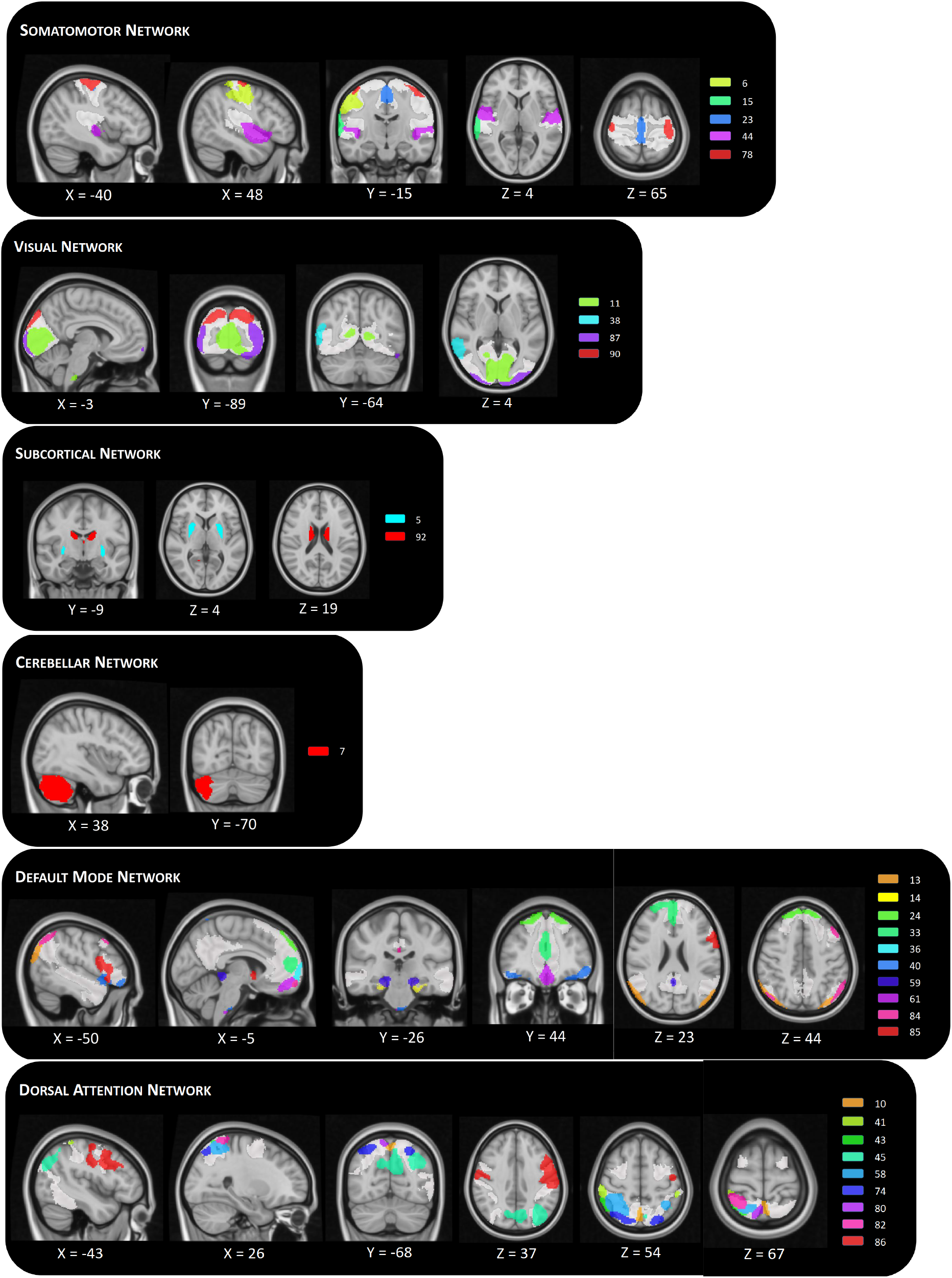
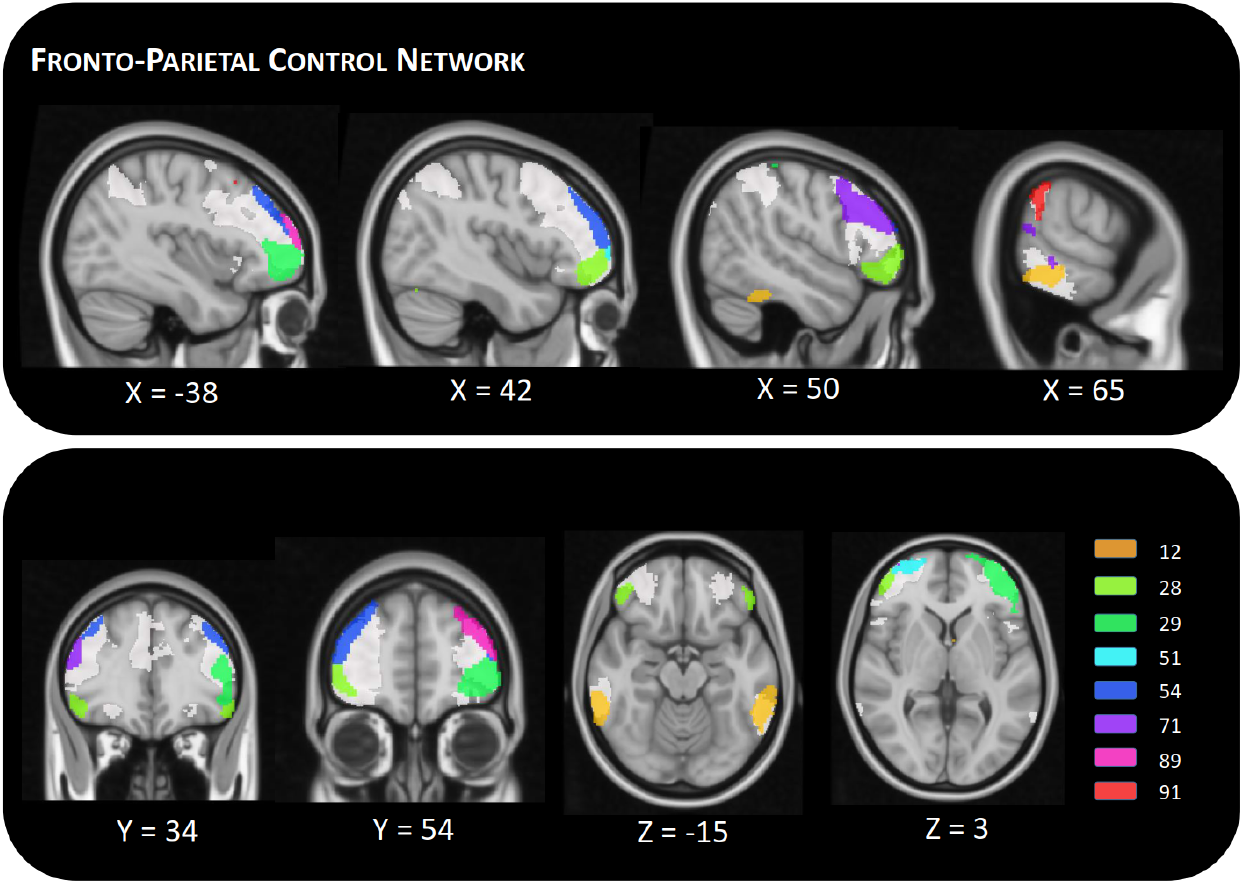
Included independent components in MNI space. Maps show the 39 identified signal components sorted into seven intrinsic functional connectivity networks according to [1], which are displayed transparent. Each color corresponds to a different component. For visualization purposes, maps show only the 60% highest values of components values. Component labels and peak coordinates are provided in supplementary table 2.

**Supplementary Fig. 2:**
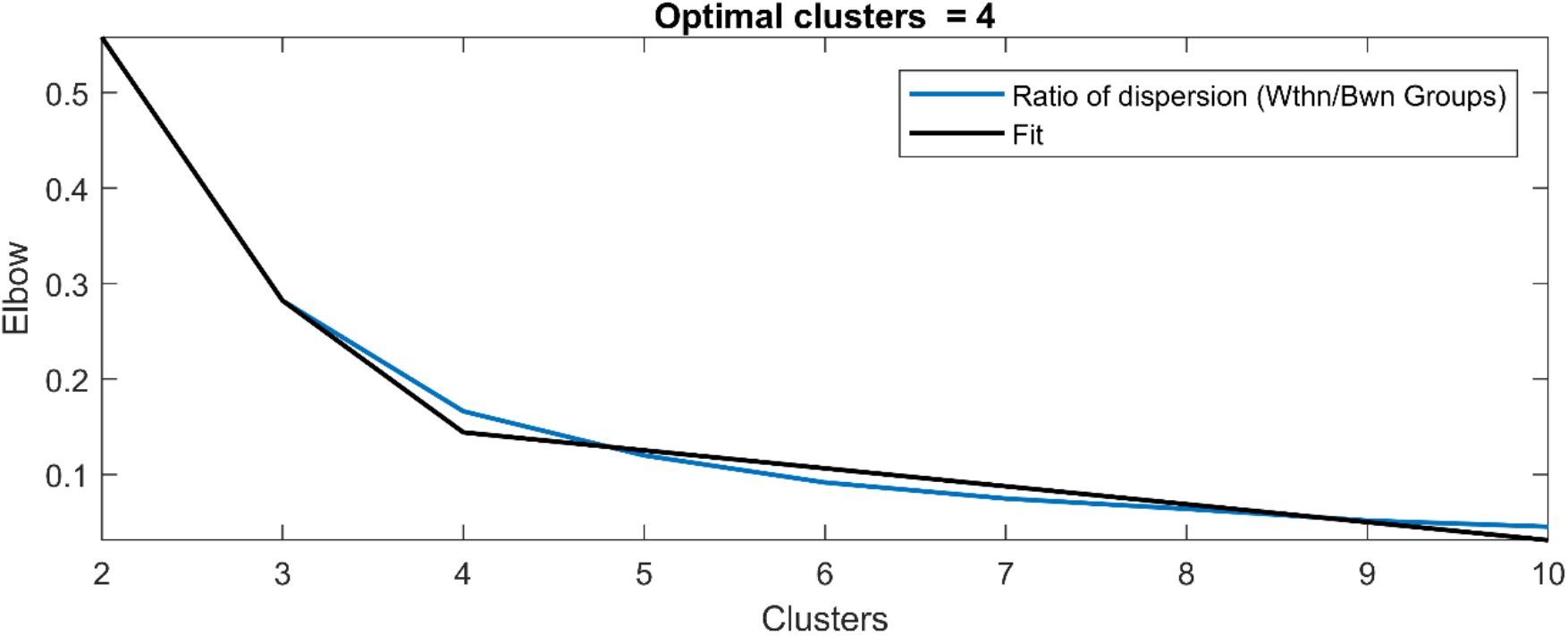
Visualization of elbow criterion.

**Supplementary Fig. 3:**
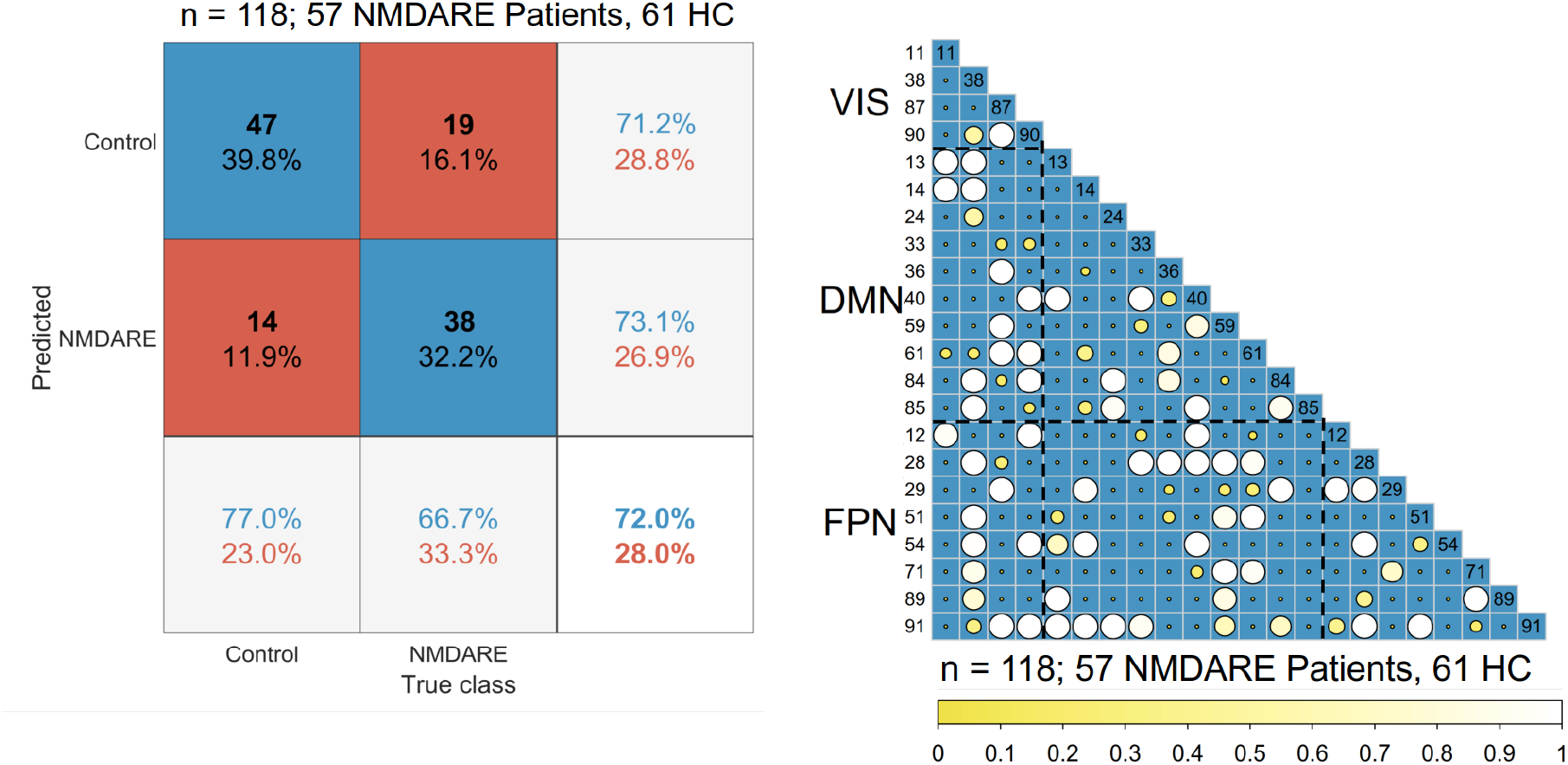
Confusion plot and feature selection matrix for static FC. Feature selection matrices showing all features that were selected for classification in at least 10% (threshold ≥ 0.1) of the classification after hyperparameter optimization (L1 regularization). Bigger and brighter circles indicate a higher selection rate (in percent/100) for classification. A key for the region numbers is provided in supplementary table 2. VIS = visual network; DMN = default mode network; FPN = fronto-parietal network; HC = healthy controls.

**Supplementary Fig. 4:**
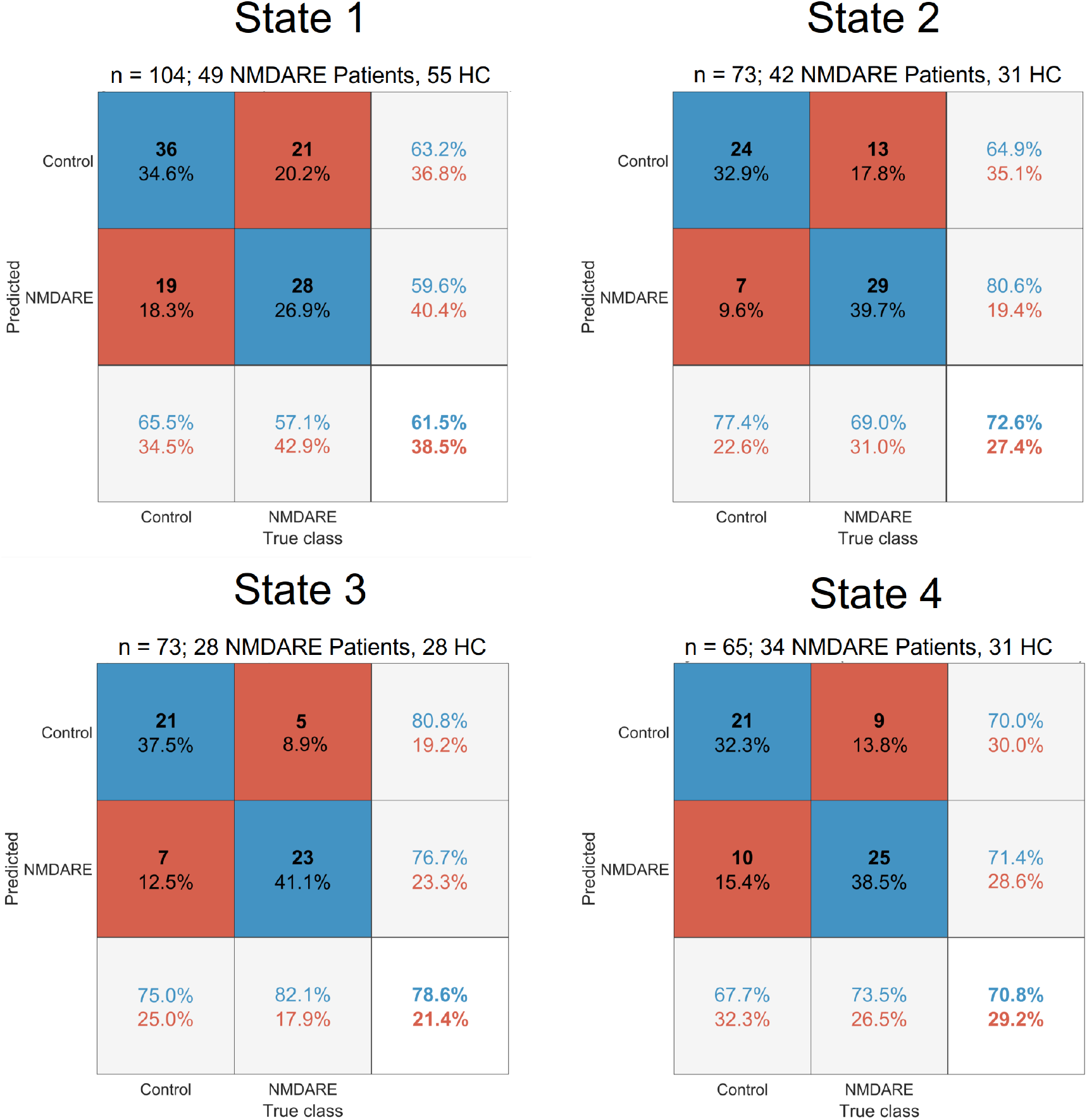
Standard confusion matrix for each state. Matrices indicate classification performance (i.e., true and false positive and negative rates and overall accuracy). HC = healthy controls.

