## Supplementary material for "State-dependent signatures of Anti-NMDA Receptor Encephalitis: a dynamic functional connectivity study"

### Supplemental material

**Supplementary Table 1:** Demographic variables and clinical measures of the participants**.** Table lists median and interquartilerange (IQR) of age, mRS at scan, and disease duration. Treatment, medication, and psychiatric symptoms during disease course were evaluated using a binary (present: 'yes' vs. absent: 'no') scale. Disease duration = acute days in hospitalization; N = number of subjects.

| | | **NMDARE Patients** | **Healthy Controls** |
|---|---|---|---|
| **N** | | 57 | 61 |
| **Sex** | ♀ / ♂ | 50/7 | 54/7 |
| **Age (years)** | Median ± IQR (N) | 25.00 ± 14.50 (57) | 26.00 ± 11.00 (61) |
| **mRS at scan** | Median ± IQR (N) | 1.00 ± 1.00 (50) | .. |
| **Disease duration** | Median ± IQR (N) | 63.00 ± 56.50 (45) | .. |
| **First-line treatment** | yes/no/unknown | 51/1/5 | .. |
| **Second-line treatment** | yes/no/unknown | 27/24/6 | .. |
| **Anticonvulsant medication** | yes/no/unknown | 39/12/6 | .. |
| **Antipsychotic medication** | yes/no/unknown | 37/15/5 | .. |
| **Positive symptoms** | yes/no/unknown | 24/27/6 | .. |
| **Negative symptoms** | yes/no/unknown | 14/37/6 | .. |

**Supplementary Table 2**: Location of included independent components. Component numbers, component labels, maximum t-value, MNI-coordinates of peak voxel and number of voxels in each component counting the voxels that contain the 60% highest values.

| | | | | |
|---|---|---|---|---|
| **Somatomotor network** | | | | |
| 6 | Postcentral gyrus (right) | 4.02 | [58 -18 48] | 2731 |
| 15 | Superior temporal gyrus (right) | 3.63 | [68 -32 10] | 1268 |
| 23 | Supplementary motor area (bil.) | 4.11 | [-2 -18 62] | 1930 |
| 44 | Superior temporal gyrus (bil.) | 3.67 | [48 0 -2] | 2814 |
| 78 | Precentral gyrus (bil.) | 4.14 | [-41 -20 63] | 751 |
| **Visual network** | | | | |
| 11 | Calcarine fissure (bil.) | 4.09 | [-10 -94 -4] | 4690 |
| 38 | Temporo-parietal-occipital junction (right) | 4.17 | [62 -46 10] | 2033 |
| 87 | Middle occipital gyrus (bil.) | 1.42 | [-40 -92 -2] | 2620 |
| 90 | Superior occipital gyrus (bil.) | 3.86 | [-22 90 34] | 1251 |
| **Subcortical network** | | | | |
| 5 | Putamen (bil.) | 4.70 | [-22 10 -12] | 2249 |
| 92 | Caudate (bil.) | 3.21 | [-12 -6 18] | 1001 |
| **Cerebellar network** | | | | |
| 7 | Cerebellum (right) | 2.77 | [46 -50 -30] | 3120 |
| **Default mode network** | | | | |
| 13 | Angular gyrus (bil.) | 4.46 | [44 -74 40] | 2222 |
| 14 | Parahippocampal gyrus (right) | 4.04 | [-23 -25 -21] | 154 |
| 24 | Dorsolateral superior frontal gyrus (right) | 5.28 | [14 46 50] | 1008 |
| 33 | Medial prefrontal cortex (bil.) | 4.37 | [- 2 62 18] | 2394 |
| 36 | Medial prefrontal cortex (bil.) | 6.34 | [-2 68 2] | 564 |
| 40 | Superior temporal gyrus (left) | 2.82 | [-54 20 -6] | 1800 |
| 59 | Hippocampus (bil.) | 3.54 | [20 -16 -16] | 1270 |
| 61 | Superior frontal gyrus, medial orb (bil.) | 3.91 | [-2 58 -12] | 882 |
| 84 | Parietal lobe, angular gyrus (bil.) | 2.69 | [-50 -60 52] | 1597 |
| 85 | Inferior frontal gyrus, opercular part (left) | 3.05 | [-62 14 18] | 1487 |
| **Dorsal attention network** | | | | |
| 10 | Parieto-occipital sulcus (bil.) | 9.20 | [-2 -60 64] | 411 |

| | | | | |
|---|---|---|---|---|
| 41 | Postcentral gyrus (left) | 5.84 | [48 -34 62] | 563 |
| 43 | Interparietal sulcus (right) | 6.13 | [44 -50 62] | 494 |
| 45 | Precuneus (bil.) | 2.99 | [-36 -74 40] | 2768 |
| 58 | Superior parietal gyrus (bil.) | 4.33 | [38 -52 60] | 2017 |
| 74 | Superior parietal gyrus (bil.) | 4.95 | [30 -68 56] | 875 |
| 80 | Parieto-occipital sulcus (right) | 6.55 | [4 -56 72] | 260 |
| 82 | Postcentral gyrus (right) | 3.89 | [28 -46 72] | 584 |
| 86 | Postcentral gyrus (bil.) | 3.46 | [-58 -6 40] | 3042 |
| **Frontoparietal network** | | | | |
| 12 | Inferior temporal gyrus (bil.) | 3.24 | [-64 -44 -14] | 1544 |
| 28 | Middle frontal gyrus, orbital part (right) | 3.67 | [44 48 -6] | 985 |
| 29 | Middle frontal gyrus, orbital part (left) | 3.43 | [-46 50 -4] | 1674 |
| 51 | Dorsolateral superior frontal gyrus (right) | 4.22 | [28 66 6] | 322 |
| 54 | Middle frontal gyrus (bil.) | 4.13 | [32 50 38] | 1533 |
| 71 | Inferior frontal gyrus, triangular part (bil.) | 3.66 | [-56 20 32] | 1800 |
| 89 | Superior frontal gyrus (left) | 5.83 | [-24 66 17] | 575 |
| 91 | Superior temporal gyrus (left) | 1.81 | [-54 20 -8] | 5643 |

**Supplementary Table 3:** Two-way ANOVA for overall connectivity. * indicates significant effect.

| | **Sum of squares** | **Df** | **F** | **p** |
|---|---|---|---|---|
| Main effect: group | 0.002 | 1 | 2.52 | 0.11 |
| Main effect: state | 0.188 | 3 | 67.62 | <0.0001 * |
| Interaction effect | 0.004 | 3 | 1.58 | 0.19 |
| Residuals | 0.268 | 290 | | |

**Supplementary Table 4**: Average windows-wise overall connectivity (± SD) across all subjects.

| | **Mean (± SD)** |
|---|---|
| State 1 | 0.23 (± 0.02) |
| State 2 | 0.27 (± 0.03) |
| State 3 | 0.30 (± 0.04) |
| State 4 | 0.24 (± 0.03) |

**Supplementary Table 5**: Post-hoc Kruskal-Wallis test to examine state-wise differences in overall connectivity ($Chi^2$=124.37, p < 0.0001, df =3). The table contains the Bonferroni-corrected p-values for pairwise state comparison.

| **State** | ***p*** |
|---|---|
| State 1 - State 2 | < 0.0001 |
| State 1 - State 3 | < 0.0001 |
| State 1 - State 4 | 0.19 |
| State 2 - State 3 | 0.11 |
| State 2 - State 4 | < 0.0001 |
| State 3 - State 4 | < 0.0001 |

**Supplementary Table 6**: Two-way ANOVA for modularity. * indicates significant effect.

| | **Sum of squares** | **Df** | **F** | **p** |
|---|---|---|---|---|
| Main effect: group | 0.012 | 1 | 3.16 | 0.076 |
| Main effect: state | 1.113 | 3 | 98.11 | <0.0001 * |
| Interaction effect | 0.002 | 3 | 0.14 | 0.94 |
| Residuals | 1.098 | 290 | | |

**Supplementary Table 7**: Average window-wise modularity (± SD) across all subjects.

| | **Mean (± SD)** |
|---|---|
| State 1 | 0.37 (± 0.06) |
| State 2 | 0.42 (± 0.07) |
| State 3 | 0.25 (± 0.05) |
| State 4 | 0.41 (± 0.07) |

**Supplementary Table 8**: Post-hoc Kruskal-Wallis test to examine state-wise differences in modularity ($Chi^2$=136.08, p < 0.0001, df =3). The table contains the Bonferroni-corrected p-values for pairwise state comparison.

| **State** | ***p*** |
|---|---|
| State 1 - State 2 | < 0.0001 |
| State 1 - State 3 | < 0.0001 |
| State 1 - State 4 | 0.0070 |
| State 2 - State 3 | < 0.0001 |
| State 2 - State 4 | 1 |
| State 3 - State 4 | < 0.0001 |

**Supplementary Table 9:** Group differences in occurrences of states**.** Group differences were calculated using the z-test for population proportions. * p < 0.05 (uncorrected).

| | **State** | **NMDARE Patients** (N, %) | **Healthy Controls** (N, %) | **z** | $p_{uncorr}$ |
|---|---|---|---|---|---|
| **Occurrence** | 1 | N=49, 85.96% | N=55, 90.16% | 0.70 | 0.48 |
| | 2 | N=42, 73.68% | N=31, 50.82% | 2.34 | 0.019* |
| | 3 | N=28, 49.12% | N=28, 45.90% | -0.35 | 0.73 |
| | 4 | N=34, 59.65% | N=31, 50.82% | -0.96 | 0.34 |

**Supplementary Table 10**: Two-way ANOVA for dwell time. * indicates significant effect.

| | **Sum of squares** | **Df** | **F** | **p** |
|---|---|---|---|---|
| Main effect: group | 23068.00 | 1 | 6.79 | 0.0096 * |
| Main effect: state | 68622.00 | 3 | 6.73 | 0.00021 * |
| Interaction effect | 20411.00 | 3 | 2.00 | 0.11 |
| Residuals | 985147.00 | 290 | | |

**Supplementary Table 11**: Two-way ANOVA for transition frequencies. * indicates significant effect.

| | **Sum of squares** | **Df** | **F** | **p** |
|---|---|---|---|---|
| Main effect: group | 7.25 | 1 | 4.07 | 0.044 * |
| Main effect: state | 46.87 | 5 | 5.26 | <0.0001 * |
| Interaction effect | 9.20 | 5 | 1.03 | 0.40 |
| Residuals | 1239.96 | 696 | | |

**Supplementary Table 12**: Two-way ANOVA for fraction time. * indicates significant effect.

| | **Sum of squares** | **Df** | **F** | **p** |
|---|---|---|---|---|
| Main effect: group | 0.036 | 1 | 0.35 | 0.56 |
| Main effect: state | 1.515 | 3 | 4.94 | 0.0023 * |
| Interaction effect | 0.037 | 3 | 0.12 | 0.95 |
| Residuals | 29.63 | 290 | | |

**Supplementary Table 13:** Differences between states in dwell time (windows), transition frequencies between states (absolute numbers), and fraction time (percentage). Differences between states were calculated using a Tukey's test. T-values and p-values are shown. * p < 0.05 (FDR-corrected), ** p < 0.01 (FDR-corrected).

| | **State** | ***t*** | $p_{FDR}$ |
|---|---|---|---|
| **Dwell time** | 1 - 2 | 3.77 | 0.0011 ** |
| | 1 - 3 | 3.61 | 0.0021 ** |
| | 1 - 4 | 1.86 | 0.25 |
| | 2 - 3 | -0.04 | 0.99 |
| | 2 - 4 | -1.69 | 0.33 |
| | 3 - 4 | -1.61 | 0.37 |
| **Transition frequency** | 1 - 2 vs 1 - 3 | 0.00 | 1.00 |
| | 1 - 2 vs 1 - 4 | -1.22 | 0.83 |
| | 1 - 2 vs 2 - 3 | 0.54 | 0.99 |
| | 1 - 2 vs 2 - 4 | 1.97 | 0.36 |
| | 1 - 2 vs 3 - 4 | 3.32 | 0.012 * |
| | 1 - 3 vs 1 - 4 | -1.22 | 0.83 |
| | 1 - 3 vs 2 - 3 | 0.54 | 0.99 |
| | 1 - 3 vs 2 - 4 | 1.97 | 0.36 |
| | 1 - 3 vs 3 - 4 | 3.32 | 0.012 * |
| | 1 - 4 vs 2 - 3 | 1.76 | 0.49 |
| | 1 - 4 vs 2 - 4 | 3.19 | 0.019 * |
| | 1 - 4 vs 3 - 4 | 4.55 | 0.00012 ** |
| | 2 - 3 vs 2 - 4 | 1.42 | 0.71 |
| | 2 - 3 vs 3 - 4 | 2.78 | 0.062 |
| | 2 - 4 vs 3 - 4 | 1.357 | 0.75 |
| **Fraction time** | 1 - 2 | 3.226 | 0.0075 ** |
| | 1 - 3 | 3.017 | 0.015 * |
| | 1 - 4 | 2.170 | 0.13 |
| | 2 - 3 | -0.093 | 0.99 |
| | 2 - 4 | -0.934 | 0.79 |
| | 3 - 4 | -0.817 | 0.85 |

**Supplementary Table 14:** Pearson's correlation coefficient between the participants' average static FC and the participants' average of each state.

| | **R** |
|---|---|
| Static – State 1 | 0.94 |
| Static – State 2 | 0.91 |
| Static – State 3 | 0.87 |
| Static – State 4 | 0.71 |

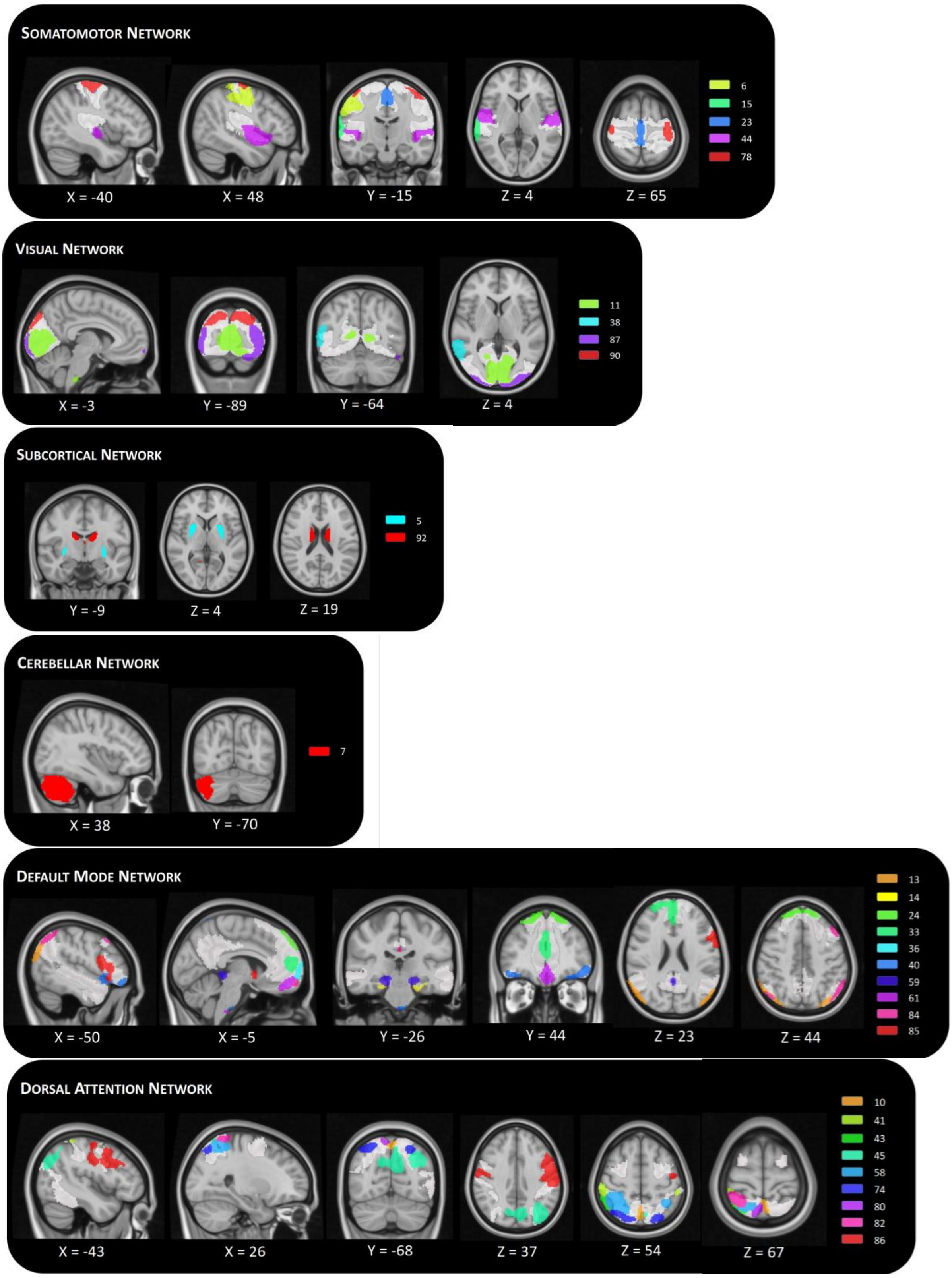
SOMATOMOTOR NETWORK
X = -40
X = 48
Y = -15
Z = 4
Z = 65
6
15
23
44
78
VISUAL NETWORK
X = -3
Y = -89
Y = -64
Z = 4
11
38
87
90
SUBCORTICAL NETWORK
Y = -9
Z = 4
Z = 19
5
92
CEREBELLAR NETWORK
X = 38
Y = -70
7
DEFAULT MODE NETWORK
X = -50
X = -5
Y = -26
Y = 44
Z = 23
Z = 44
13
14
24
33
36
40
59
61
84
85
DORSAL ATTENTION NETWORK
X = -43
X = 26
Y = -68
Z = 37
Z = 54
Z = 67
10
41
43
45
58
74
80
82
86

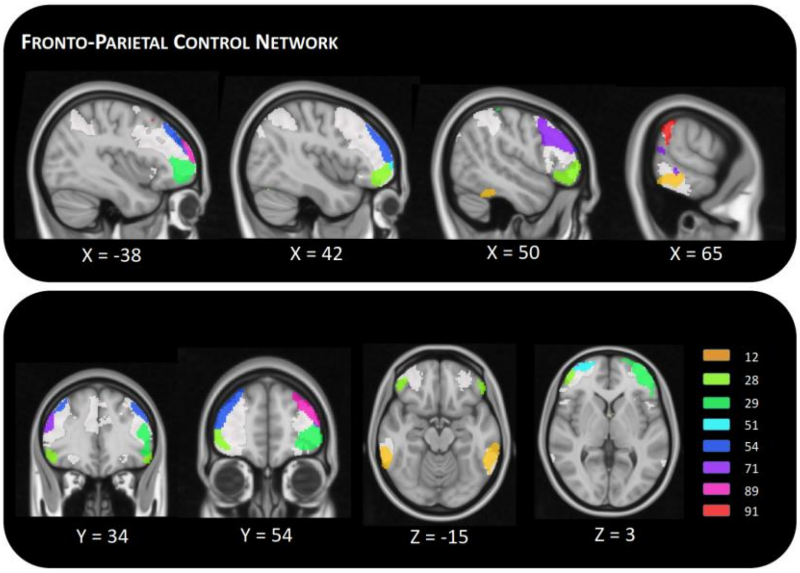


**Supplementary Fig. 1:** Included independent components in MNI space. Maps show the 39 identified signal components sorted into seven intrinsic functional connectivity networks according to [1], which are displayed transparent. Each color corresponds to a different component. For visualization purposes, maps show only the 60% highest values of components values. Component labels and peak coordinates are provided in Supplementary Table 2.

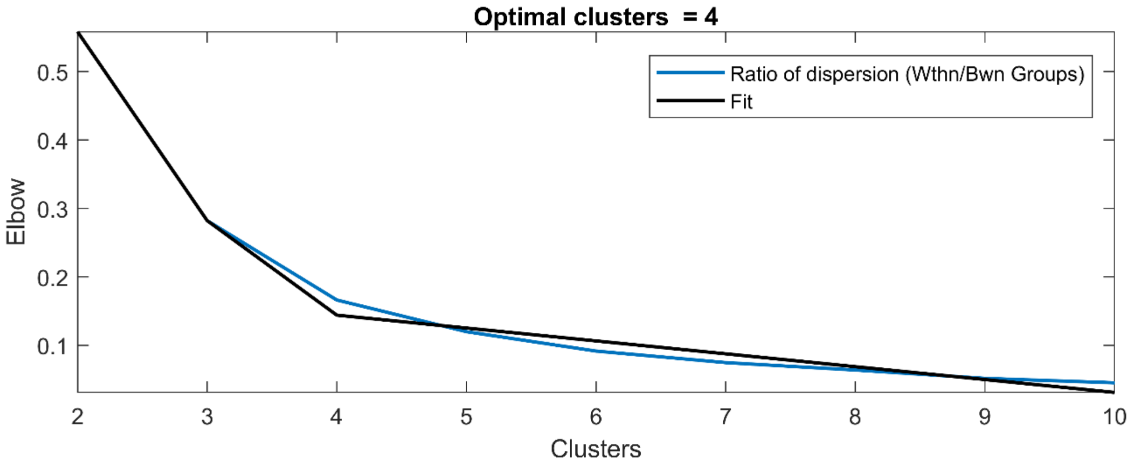


**Supplementary Fig. 2:** Visualization of elbow criterion.

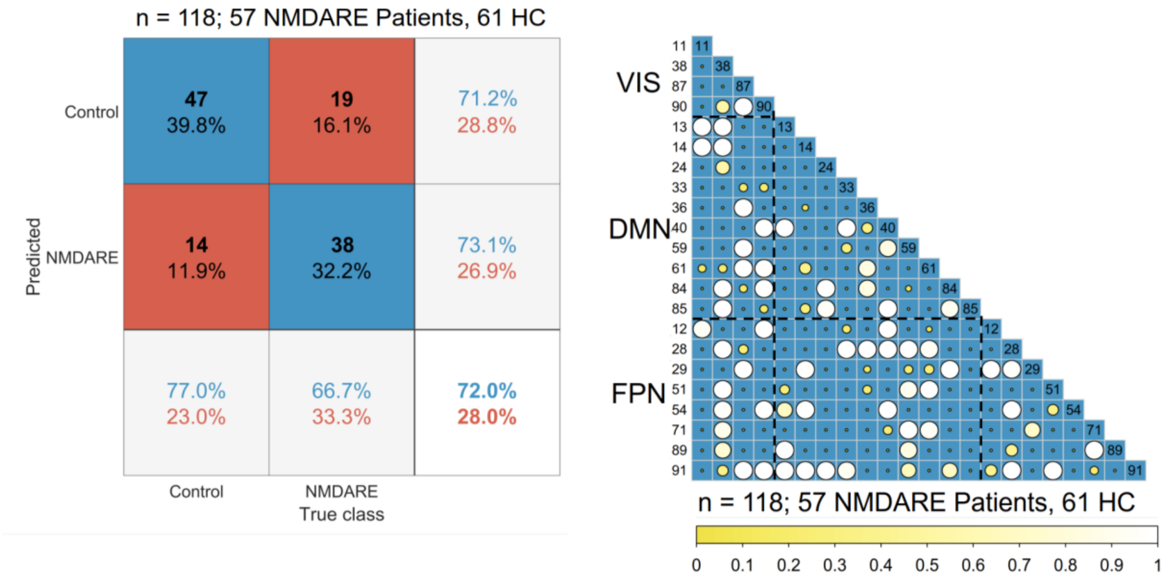


**Supplementary Fig. 3:** Confusion plot and feature selection matrix for static FC. Feature selection matrices showing all features that were selected for classification in at least 10% (threshold ≥ 0.1) of the classification after hyperparameter optimization (L1 regularization). Bigger and brighter circles indicate a higher selection rate (in percent/100) for classification. A key for the region numbers is provided in Supplementary Table 2. VIS = visual network; DMN = default mode network; FPN = fronto-parietal network; HC = healthy controls.

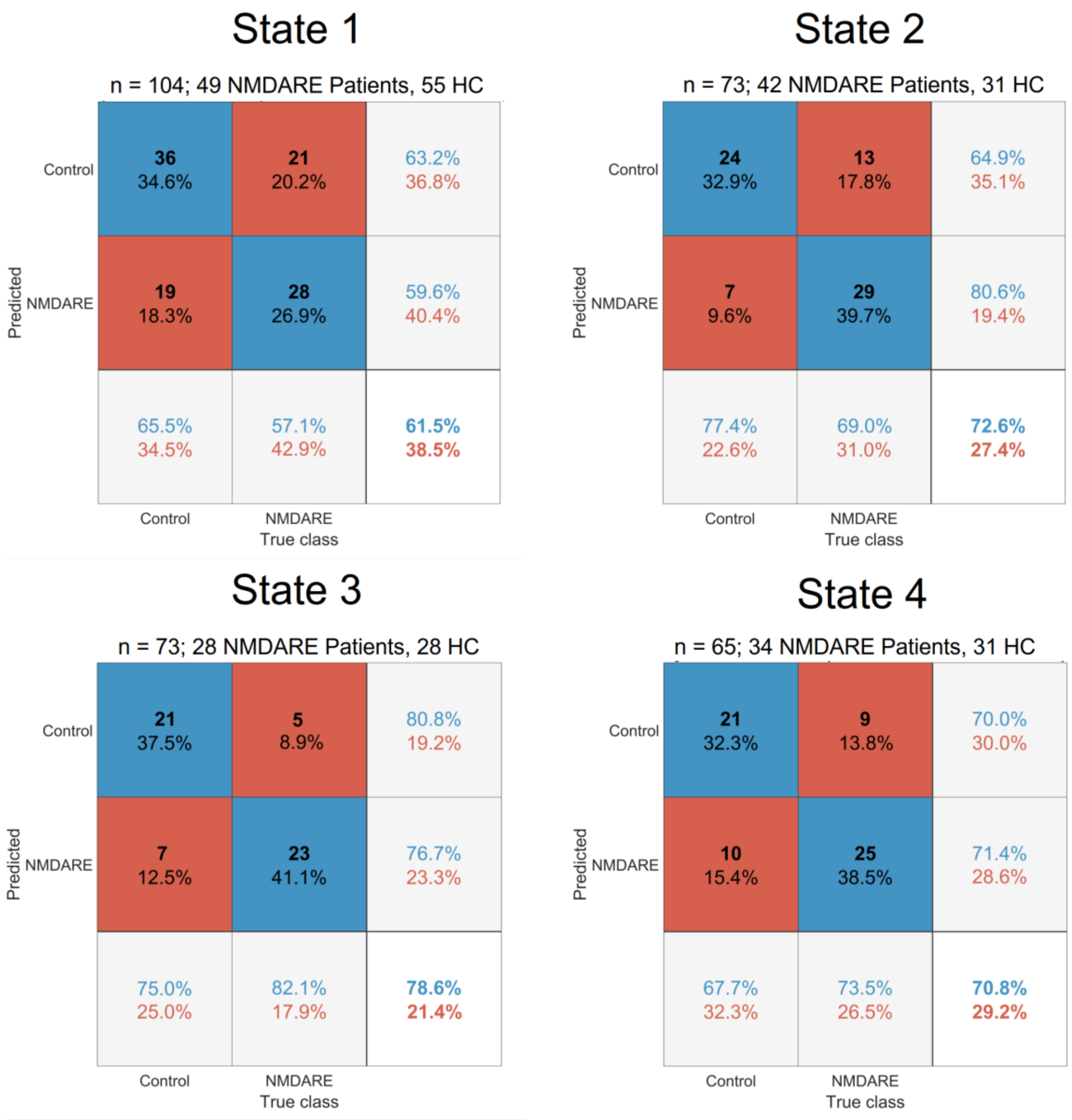


**Supplementary Fig. 4:** Standard confusion matrix for each state. Matrices indicate classification performance (i.e., true and false positive and negative rates and overall accuracy). HC = healthy controls.
